# Systematic assessment of the impact of targeted selection methods and environment-mimicking culture conditions on fungal natural product libraries

**DOI:** 10.64898/2026.03.04.709592

**Authors:** Monica Ness, Karen Wendt, Thilini Peramuna, Demetrius I. Tillery, Jillian E. Murray, Robert H Cichewicz, Laura-Isobel McCall

**Affiliations:** Department of Chemistry and Biochemistry, University of Oklahoma, Norman, Oklahoma 73019, United States; Department of Chemistry and Biochemistry, San Diego State University, San Diego, California, 92182, United States; College of Pharmacy, University of Michigan, Ann Arbor, Michigan, 48109, United States

**Keywords:** Natural Products, Fungi, Extract libraries, OSMAC, Mass Spectrometry

## Abstract

Natural products are a rich source of bioactive molecules and undiscovered chemical scaffolds with significant potential for novel drug discovery. Among these, fungi are particularly promising, offering diverse metabolites and undiscovered structural motifs. Large, well-curated collections of crude extracts, or “libraries”, are central to fungal natural product discovery, serving as starting material for bioassay-guided isolation of new compounds. However, the systematic influence of fungal selection strategies, culturing methods, and environmental factors on chemical diversity remains underexplored. In this study, we analyzed several large fungal libraries to assess how geographic origin, and phylogenetic classification shape fungal chemical profiles. We also evaluated whether culturing conditions that more closely mimic natural environments can enhance metabolite diversity. Our findings offer practical guidelines for optimizing fungal natural product library design, improving drug development efficiency and access to novel chemotypes for future drug discovery.

**Summary Figure:** 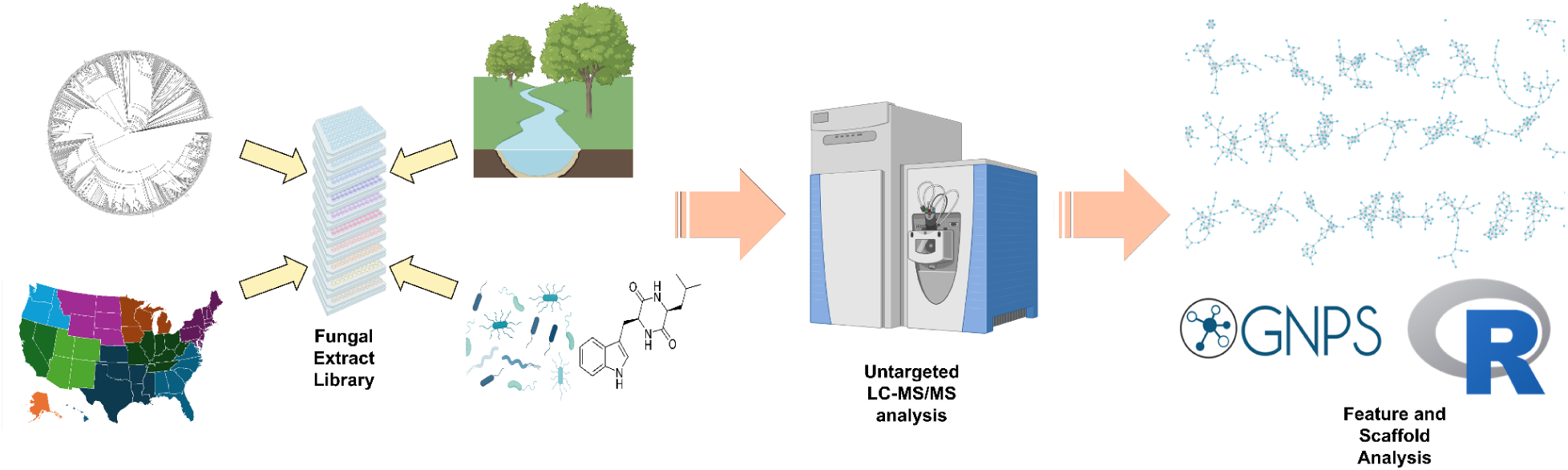

## Introduction

Natural products (NPs) remain one of the most important sources of novel bioactive molecules, particularly in drug discovery. Fungal natural products are of special interest due to their chemical diversity and complex ecological roles, yet efforts to systematically mine fungi for new metabolites face longstanding challenges. A central bottleneck lies in the curation of crude extract libraries. Large libraries are difficult and expensive to maintain, and as collections expand, the rate of novel discoveries becomes more time-consuming. Bioactivity-based screening discovery thus becomes a “needle in a haystack” problem, with diminishing returns as libraries grow ^1,2^ and with risk of frequent rediscovery of known NPs, wasting time and resources.

Despite the importance of library curation, few systematic studies on how to properly build a new library have been done ^3^. There are multiple studies that implement a “One Strain Many Compounds (“OSMAC”) approach, where culturing conditions are varied to induce expression of cryptic biosynthetic gene clusters. Importantly, rather than broadly assessing how OSMAC methods impacted the overall chemical profile of their collections ^4,5^, most of these studies emphasize one or two novel chemotypes ^6,7^, highlighting individual success stories. This positive-results bias obscures the true effectiveness of different collection and culturing methods, leaving gaps in guidance for researchers aiming to build efficient, diverse libraries.

To maximize the chemical space represented in libraries, a variety of strategies have been employed. Some groups emphasize regional or geographic sampling, assuming that distinct climates or habitats yield unique chemistry ^8,9^. Others turn to phylogenetic approaches, using sequence-based distances to guide inclusion of multiple strains ^10,11^. On the cultivation side, numerous methods have been explored, ranging from altering nutrient composition to adding environmental signals or microbial cofactors ^12,13^. Again, there is little reporting of overall impact on the crude extract collection profile with multiple genera.

In this work, we sought to address these gaps through a systematic analysis.

Using untargeted LC-MS/MS, feature-based molecular networking ^14^, and accumulation curves compared to random selection baselines, we evaluated the impact of selection methods (geographic and phylogenetic) and environmental mimicry approaches (soil extract, lipopolysaccharide, and diketopiperazine supplementation).

Our results show that selection strategies have surprisingly little impact on scaffold accumulation, contrary to intuitive expectations. In contrast, culturing conditions produced varied effects, with certain interventions transiently suppressing diversity and others enhancing it in a genus-or time-dependent manner. By systematically comparing these approaches, we aim to provide clarity for fungal library design moving forward, highlighting practical strategies for maximizing scaffold-level chemical diversity while avoiding inefficient or ineffective practices.

## Results

### Effect of Sample Selection Methods

To start any new crude extract library, one must consider how to select the extracts for the collection. Ideally, any choice made would maximize the chemical diversity of the entire collection while reducing repetitiveness, and, at the same time, minimize the amount of initial work needed to collect the samples. In the first part of our meta-analysis, we analyze the two selection methods for fungal extract collection, particularly geographic and phylogeny-based methods, all of which are compared to random selection.

We apply these methods to a collection of extracts from the genus *Trichoderma*. The *Trichoderma* collection, which contains 835 samples, representing all NOAA geographic regions in the United States. This collection was analyzed via untargeted LC-MS/MS. Notably, the *Trichoderma* isolates constitute a diverse dataset, where over 65,000 features and 9500 scaffolds were detected. For this analysis, we focused on scaffold diversity determined with GNPS Feature-Based Molecular Networking ^14^. When doing the same analysis on the feature levels, we saw similar results. Those can be seen in Figures S1-S3 and Tables S1-S2

To see if the same results apply to a broader collection, we also re-analyzed untargeted LC-MS/MS data on a dataset of 1439 fungal extracts, which contains 141 different genera. This collection is also very diverse, containing samples from each US state, and totals over 4000 scaffolds and 35,000 features. This collection is called the multi-genera collection.

### Geography-based Selection

A common assumption in natural products research is that maximizing the geographic spread of samples will maximize chemical diversity, since different climates and environments are expected to drive evolution of unique metabolites ^8,9^. Several studies emphasize wide geographic range strategies, where collections are designed to cover as many regions as possible, with the assumption that it can increase the likelihood of discovering novel chemical scaffolds ^15,16^. Others explicitly highlight broad regional sampling as a strength of their extract libraries, equating diversity of region with chemical diversity ^17^. In our analysis, we divided samples by region of origin according to NOAA climate zone definition ^18^. Scaffold diversity determined via Feature-Based Molecular Networking ^14^.

To understand the effectiveness of a geographically spread library, we simulated a scenario where samples were chosen based on maximizing regional spread according to NOAA climate regions, starting at a random point each time. Twenty-five different iterations were performed in the analysis and compared to random selections (Fig. 1). We found that prioritizing regional diversity does not promote more rapid library diversity. This indicates that prioritizing geographic distribution does not translate into faster accumulation of chemical diversity with fewer samples, as intentionally maximizing regional diversity did not outperform random sample selection for inclusion in the library.

**Figure 1.**
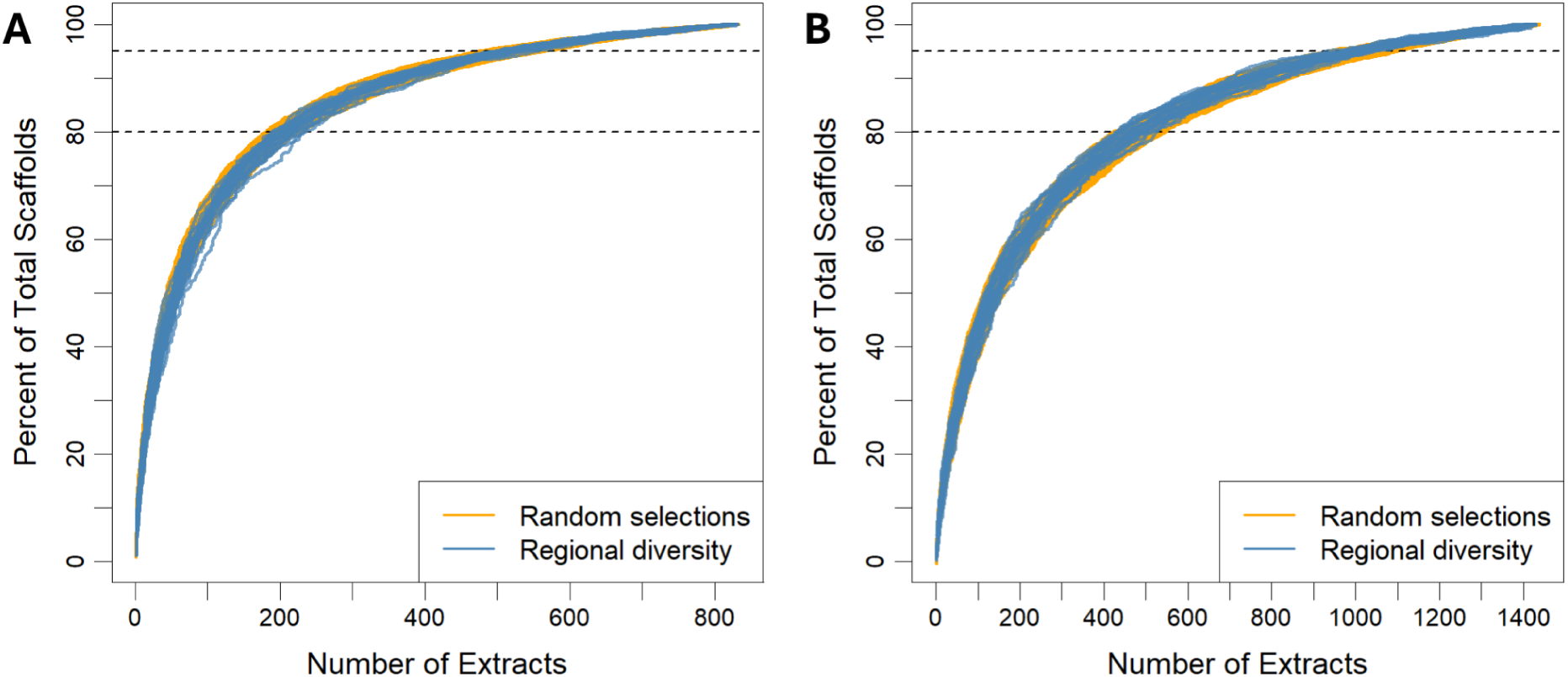
Regional diversity prioritization does not outperform random selection. We simulated a geographic diversity prioritization in our *Trichoderma* (A) and multi-genera collection (B) library, where each extract added to the library was intended to maximize regional scope. Twenty-five iterations of this were done and compared to 25 iterations of random selection, with no improvement in the rate of accumulation of chemical diversity.

When average scaffold counts per sample were calculated, most regions contributed similar levels of diversity, and substantial scaffold overlap was observed across regions (Fig. 2). Further, when analyzing distributions of unique versus shared scaffolds (Table 1, Table S3), several regions produced relatively few unique scaffolds despite extensive sampling. An initial comparison suggested that samples from southern regions contributed a larger number of unique scaffolds, potentially reflecting wetter and/or warmer climates may be more permissive for increasing fungal metabolite diversification. Consequently, concentrated sampling within certain southern regions would yield more unique chemistry. Taken together, these results suggest that while some unique scaffolds are region-specific, overall chemical diversity is driven more efficiently by concentrated sampling in productive geographies rather than maximizing coverage across multiple regions. This aligns with observations that even within a single region, sampling across different ecological niches or microclimates (e.g., different cities) can capture substantial diversity ^19^, while requiring less work on behalf of the collector.

**Figure 2.**
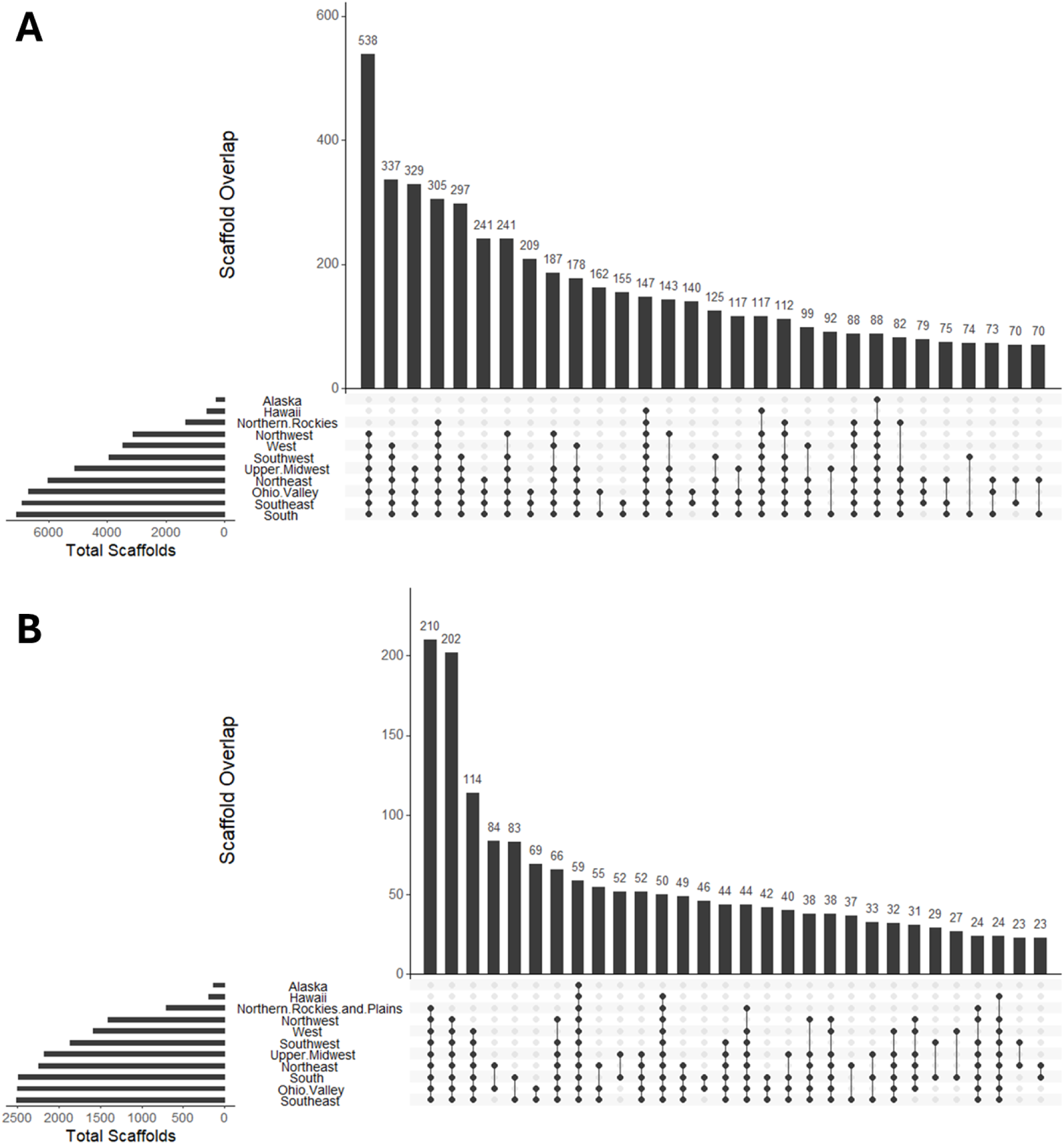
Broad overlap of shared chemical scaffolds across regions. UpSet plots showing the distribution of scaffolds shared among geographic regions. Only the top 30 intersection sets are displayed, and scaffolds detected in a single region only are excluded. The top panel shows results for *Trichoderma*, while the bottom panel shows the multi-genera dataset.

**Table 1.** Summary of average unique and total scaffolds per each region. This table shows results for the *Trichoderma* dataset. Similar patterns were seen in the muti-genera collection (Table S3).

| Region | Total Extracts | Total Scaffolds | Average Number of Scaffolds | Scaffolds Unique to Region | Average Unique Scaffolds per Extract |
| --- | --- | --- | --- | --- | --- |
| Alaska | 4 | 291 | 113 | 0 | 0.0 |
| Hawaii | 2 | 602 | 342 | 1 | 0.5 |
| Northern Rockies | 7 | 1335 | 285 | 4 | 0.6 |
| Northwest | 41 | 3157 | 214 | 41 | 1.0 |
| West | 48 | 3534 | 233 | 73 | 1.5 |
| Southwest | 35 | 4044 | 301 | 87 | 2.5 |
| Upper Midwest | 70 | 5241 | 298 | 124 | 1.8 |
| Northeast | 135 | 6126 | 296 | 91 | 0.7 |
| Ohio Valley | 137 | 6845 | 329 | 157 | 1.1 |
| Southeast | 194 | 7137 | 281 | 238 | 1.2 |
| South | 160 | 7368 | 333 | 290 | 1.8 |

### Phylogeny-based Selection

An intuitive assumption in natural product library design is that sampling across a wide range of genera and species will yield greater chemical diversity. To test this hypothesis, we examined whether phylogeny-guided selection strategies could outperform random sampling in terms of scaffold diversity.

For the *Trichoderma* collection, we used ITS and tef sequencing data to capture both broad and fine scale phylogenetic differences among isolates. (ITS: internal transcribed spacer region, commonly used for fungal genera-level identification. tef: translation elongation factor 1α, often providing higher resolution within genera ^20,21^). We then simulated a rational selection library design scenario, in which new isolates were selected to maximize phylogenetic distance from those already included in the library (See Methods: Data Processing). This was compared directly to random sampling of the same number of isolates.

Contrary to expectations, scaffold diversity did not accumulate more rapidly under the rational phylogenetic selection scheme compared to random sampling. In fact, accumulation curves showed nearly identical trajectories between the two approaches. To confirm these findings under a broader phylogenetic range, we repeated the analysis using a multi-genera collection representing a broad range of fungal genera. Here, only ITS sequencing was available and used to generate phylogenetic distance matrices. Once again, rational phylogenetic selections performed no better, and in fact worse, than random selection (Fig. 3).

**Figure 3.**
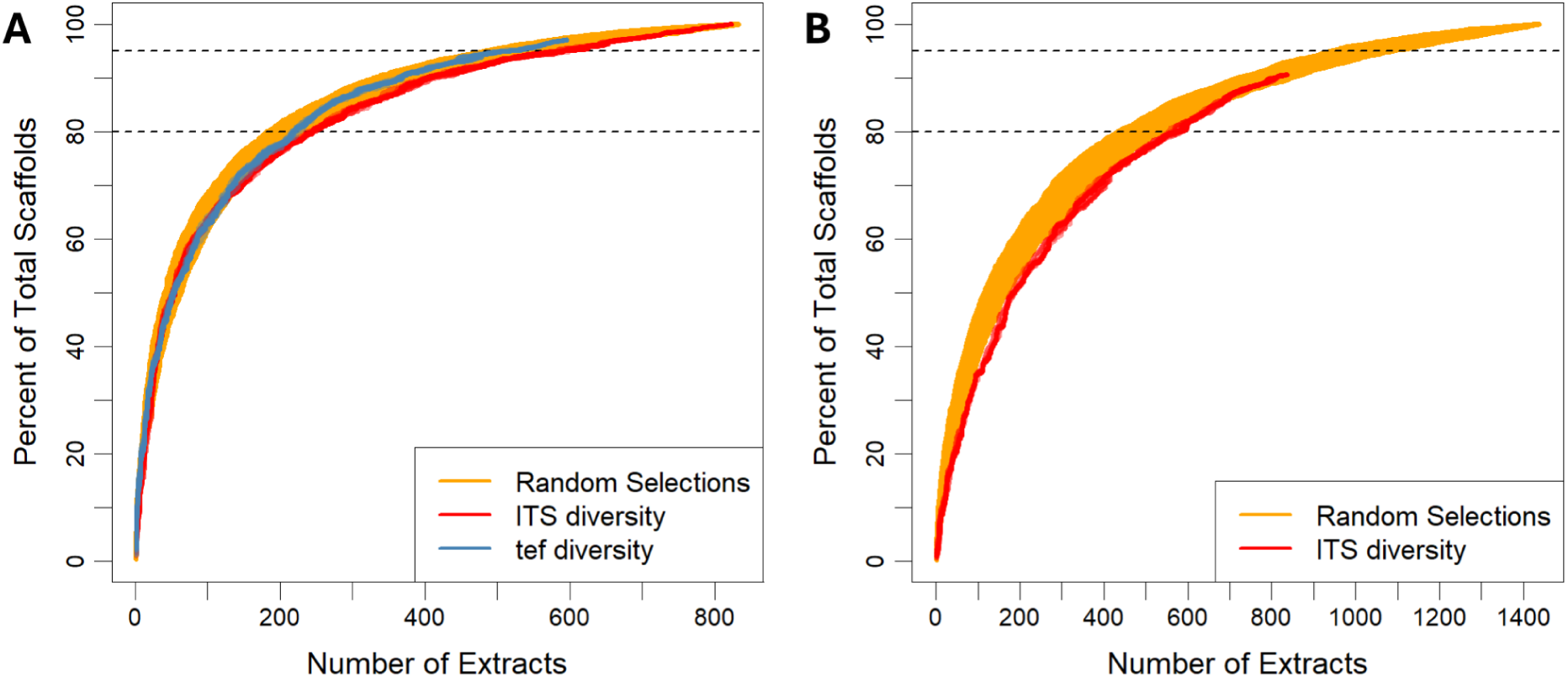
Phylogeny-based selection does not outperform random selection. Each rational phylogeny-based selection started with a random isolate, and subsequent isolates were selected to maximize phylogenetic diversity. The results for the *Trichoderma* collection are shown on the left, and the multi-genera collection is shown on the right. For both datasets, 25 iterations were used.

These results indicate that, at least for the wider and narrow phylogenetic libraries tested, maximizing phylogenetic divergence does not necessarily maximize chemical scaffold diversity. Furthermore, such an outcome draws attention to the value of testing basic presumptions that tend to drive collection strategies in favor of implementing quantitative assessment measurements that can help reduce the inefficient allocation of limited library development resources.

### Effect of Culturing Methods

The cultivation of fungal isolates following collection is a critical step in natural product discovery. While the fundamental objective of culturing is straightforward, laboratories often employ a broad range of lab-specific approaches ^22^, resulting in a lack of standardized methodology. This variability is relevant in the context of maximizing metabolite diversity, where subtle changes in culture conditions may influence chemical scaffold output.

Previous studies have suggested that the addition of exogenous factors may alter microbial behavior, potentially stimulating cryptic biosynthetic pathways ^23^.

However, the extent to which these interventions can be generalized across taxa to enhance the diversity of metabolites remains poorly characterized. Addressing this knowledge gap is especially relevant for large-scale efforts in natural product research, where broad strategies for increasing scaffold diversity are more practical than species-specific optimization.

In the present work, we examined 84 fungal isolates from three genera of fungi (*Chaetomium*, *Clonostachys*, *Purpureocillium*). Each was subjected to a consistent extraction process but cultured under multiple conditions designed to mimic elements of their natural environments. In addition, two distinct time points were evaluated to capture time-specific changes in metabolite production. Untargeted LC-MS/MS was employed to profile the resulting metabolomes, with particular emphasis on scaffold-level diversity. To see equivalent analysis on the feature-level, see Figures S4-S7, which draw the same conclusions as the scaffold-level analysis.

We tested 3 different additions to the base culturing media. First, an aqueous extract of local soil. Secondly, lipopolysaccharide, a key component of bacterial membranes. And third, different diketopiperazines. Diketopiperazines have been linked to quorum sensing and bioactivity in many natural products ^24^. Our goal was not to identify optimal conditions for each individual isolate or genus, but rather to evaluate whether a broadly applicable culturing strategy could consistently increase metabolite diversity across genera and species. Particularly, we want to test the “One Strain Many Compounds (OSMAC)” approach ^4,25^, to a large diverse fungal collection in an unbiased fashion, rather than highlight one or two success stories.

### Soil extract

One method for supplementing fungal cultures is with soil extract, with the rationale that it mimics natural environments, where many fungal species naturally grow and interact with the environment. To test this idea, we prepared an aqueous extract of soil and incorporated it into culture media under varying conditions (See Methods).

Three treatment groups were established: (1) sucrose supplementation only, (2) soil extract only, and (3) a combined sucrose-soil extract mixture. Each was evaluated at two time points: 7 and 14 days.

At the 7-day time point, supplementation had a clear impact on metabolite output.

Cultures receiving soil extract alone produced the highest scaffold counts across all three genera, while sucrose-only and combined treatments yielded lower diversity. These results suggest that the soil extract can induce early activation of biosynthetic pathways, providing an immediate but transient enhancement of chemical output (Fig. 4).

**Figure 4.**
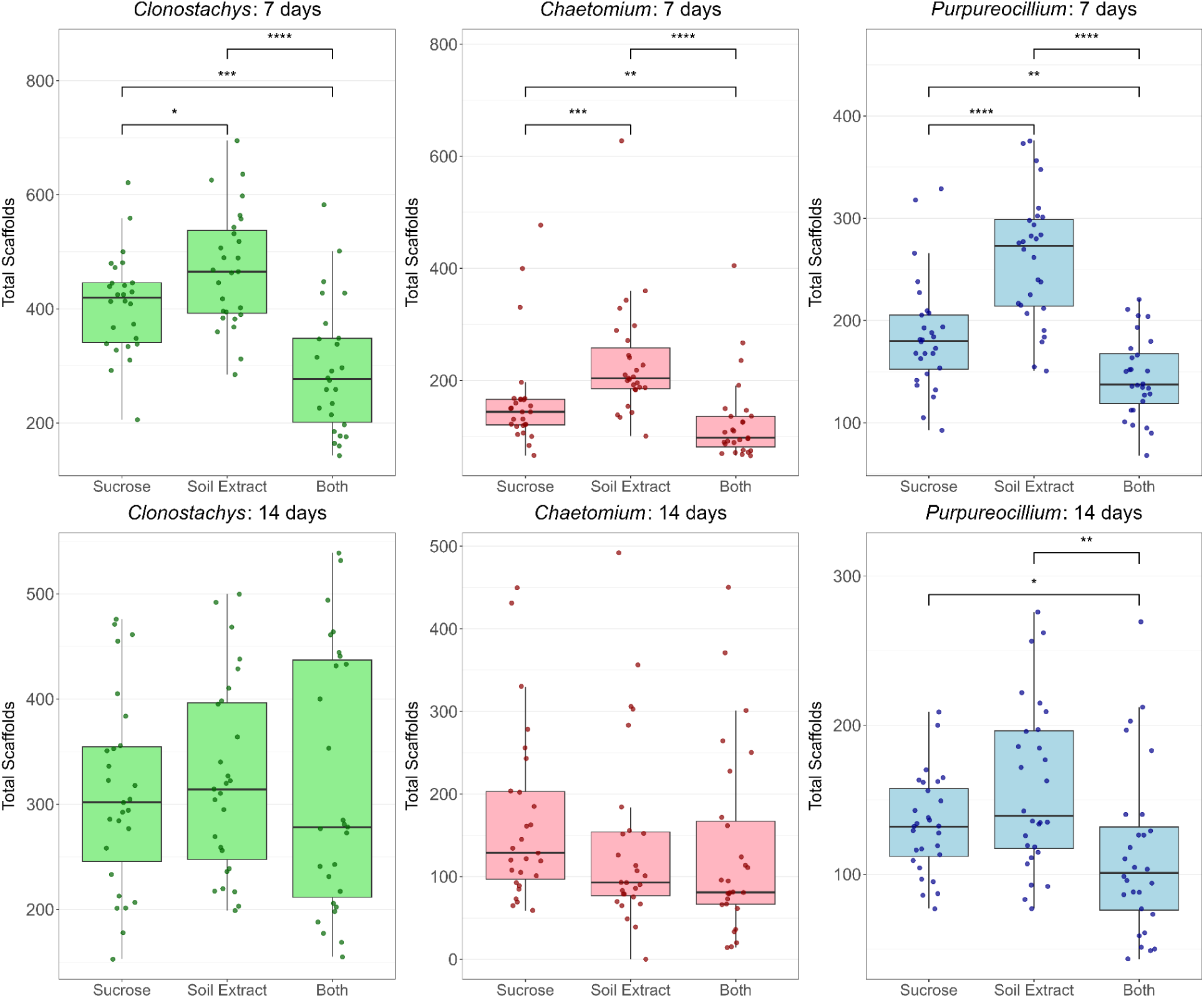
Summaries of total scaffolds per isolate, separated by genera and timepoint. Asterisks indicate statistical significance determined via Wilcoxon rank-sum test. * p < 0.05, ** p < 0.01, *** p < 0.001, **** p < 0.0001.

By the 14-day time point, scaffold-level differences between treatment groups largely disappeared, indicating that the stimulatory effect of soil extract diminishes over time. An exception was observed in *Purpureocillium*, where significant differences persisted, suggesting that some genera may maintain a longer-term response to environmental cues.

Overall, these findings support the idea that soil extract supplementation can temporarily boost chemical diversity in fungal cultures, particularly during early growth stages. However, the effect may not be universally sustained, implying that soil-derived signals may act as short-term metabolic triggers.

### Lipopolysaccharide addition

Fungi and bacteria often display a complex relationship in nature, promoting the formation of new natural products from the fungi, or inhibiting biosynthetic gene clusters^26^. In an effort to replicate environmental conditions in fungal cultures, we wanted to see if the addition of lipopolysaccharide (LPS) would stimulate metabolite production in our broad collection of fungal isolates. LPS is the key component of Gram-negative bacterial membranes, and contributes nearly 80% of the surface area of these membranes ^27^. The hypothesis behind this addition was that interaction with the LPS would be sufficient to stimulate production, without necessitating fungal-bacterial co-cultures. For this study, we used three different concentrations of LPS: 0.06, 0.6, and 1.8 ng/ml of LPS.

Similar to the soil extract culturing experiment, the effects of LPS were most dramatic at the 7 day time point. However, rather than increasing scaffold diversity, LPS addition generally resulted in a decrease in overall scaffold counts across all genera, and this effect did not appear to be proportional to concentration of LPS added, except in the *Purpureocillium* isolates. By day 14, differences between treatment groups had diminished (Fig. 5).

**Figure 5.**
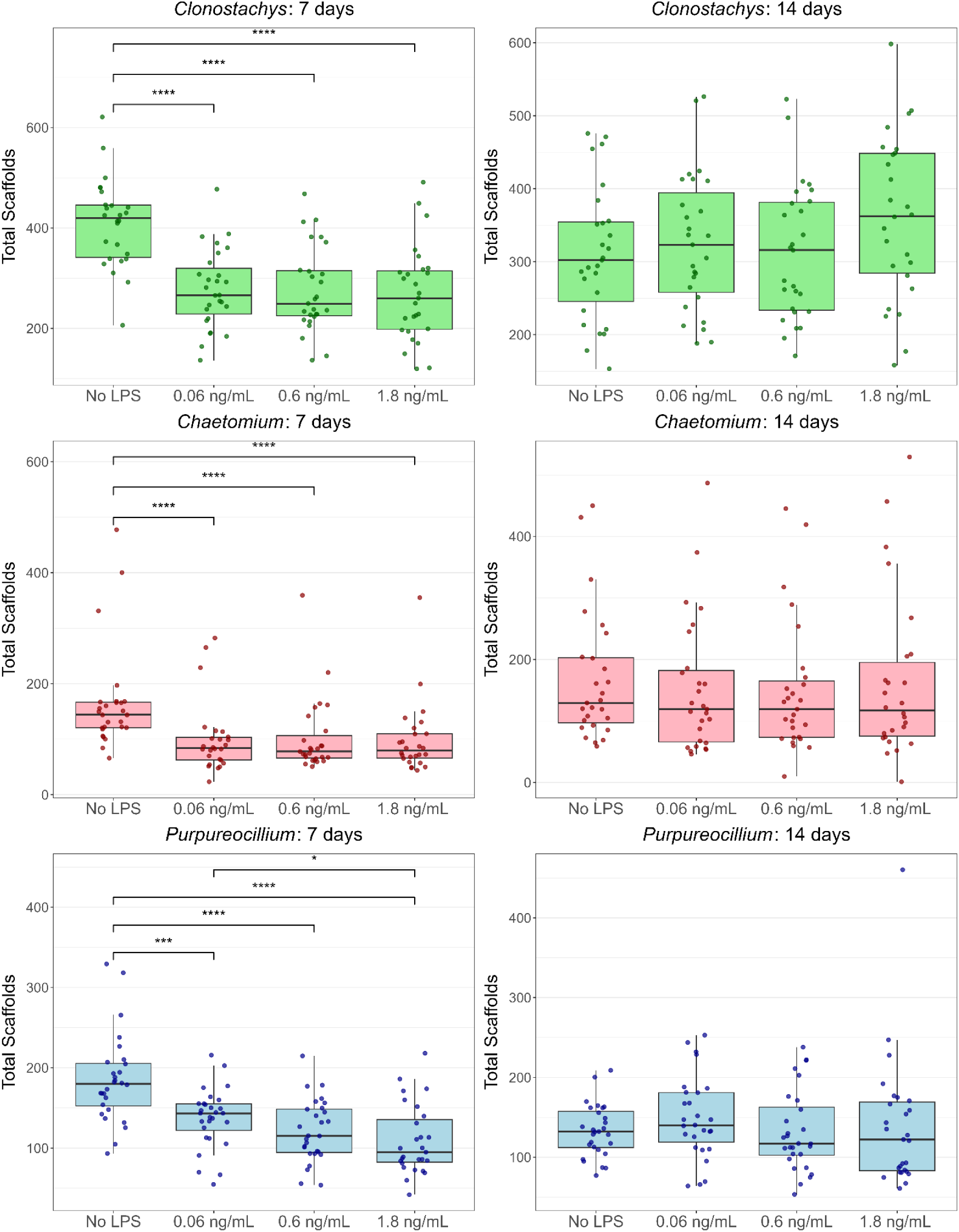
Boxplots showing scaffolds per isolate for each genera, timepoint, and LPS experimental group. Asterisks indicate statistical significance determined via Wilcoxon rank-sum test. * p < 0.05, ** p < 0.01, *** p < 0.001, **** p < 0.0001.

To investigate whether the changes represented activation of new metabolite pathways or merely diminished the existing metabolites, we compared scaffold-level overlap across treatments. While LPS supplementation did lead to the production of some new scaffolds, the majority of scaffolds overlapped with those observed in control cultures and other scaffolds were lost (Fig. 6). Treatment also changed metabolite feature abundance; however, there was an overlap in metabolite features impacted by 7-day 1.8 ng LPS treatment, those impacted by 14-day 1.8 ng LPS treatment, and those impacted by culture time alone (comparing 7-day and 14-day untreated cultures).

**Figure 6.**
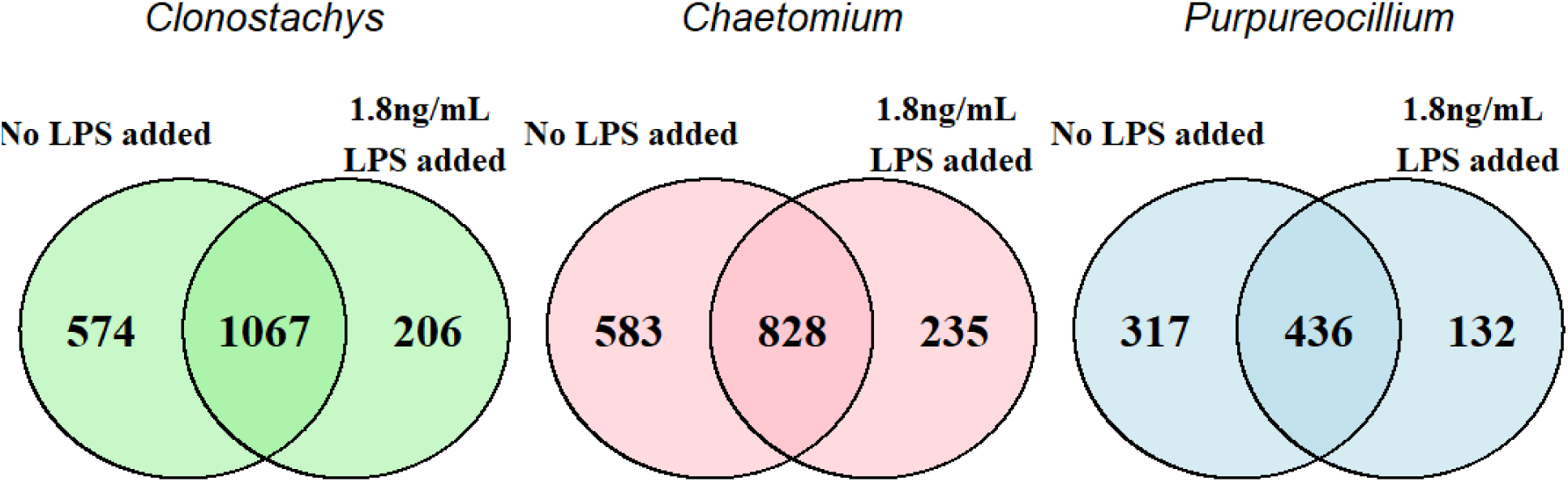
Venn diagrams showing scaffold overlaps between control and LPS groups. This data shows the 7-day timepoint.

Strikingly, the biggest overlap was observed between metabolite features impacted by 7 days of LPS treatment and those impacted by culture time, suggesting that some effects of culture additives could just be obtained by modifying culture duration instead (Fig. S9). Analysis of the 7-day and 14-day LPS treatments indicated that fatty amides were affected across all three genera. *Chaetomium* and *Purpureocillium* exhibited greater overlap in terms of chemical family annotations across both time points, with LPS treatment impacting nucleosides in these genera (Fig. S10).

Overall, these results indicate that LPS addition has a measurable impact on fungal metabolomes. It can trigger the production of specific metabolites while inhibiting the production of others, with the balance between induction and suppression depending on the timepoint and the specific fungus (Fig. S9). However, it does not broadly enhance scaffold diversity and may even suppress total scaffold output in the short term.

### Diketopiperazine addition

Diketopiperazines (DKPs) are a class of cyclic dipeptides widely linked to quorum sensing molecules in microbial systems and possess a wide range of bioactive properties, including as potential therapies to humans and stimulators and inhibitors of natural product production ^24^, making them attractive candidates for stimulating fungal metabolite production. To test their roles in fungal natural product production, we synthesized five distinct DKPs (Table 2, see Supplementary Data 1) and added them to fungal cultures at two concentrations (100 nM and 10 μM).

**Table 2.**
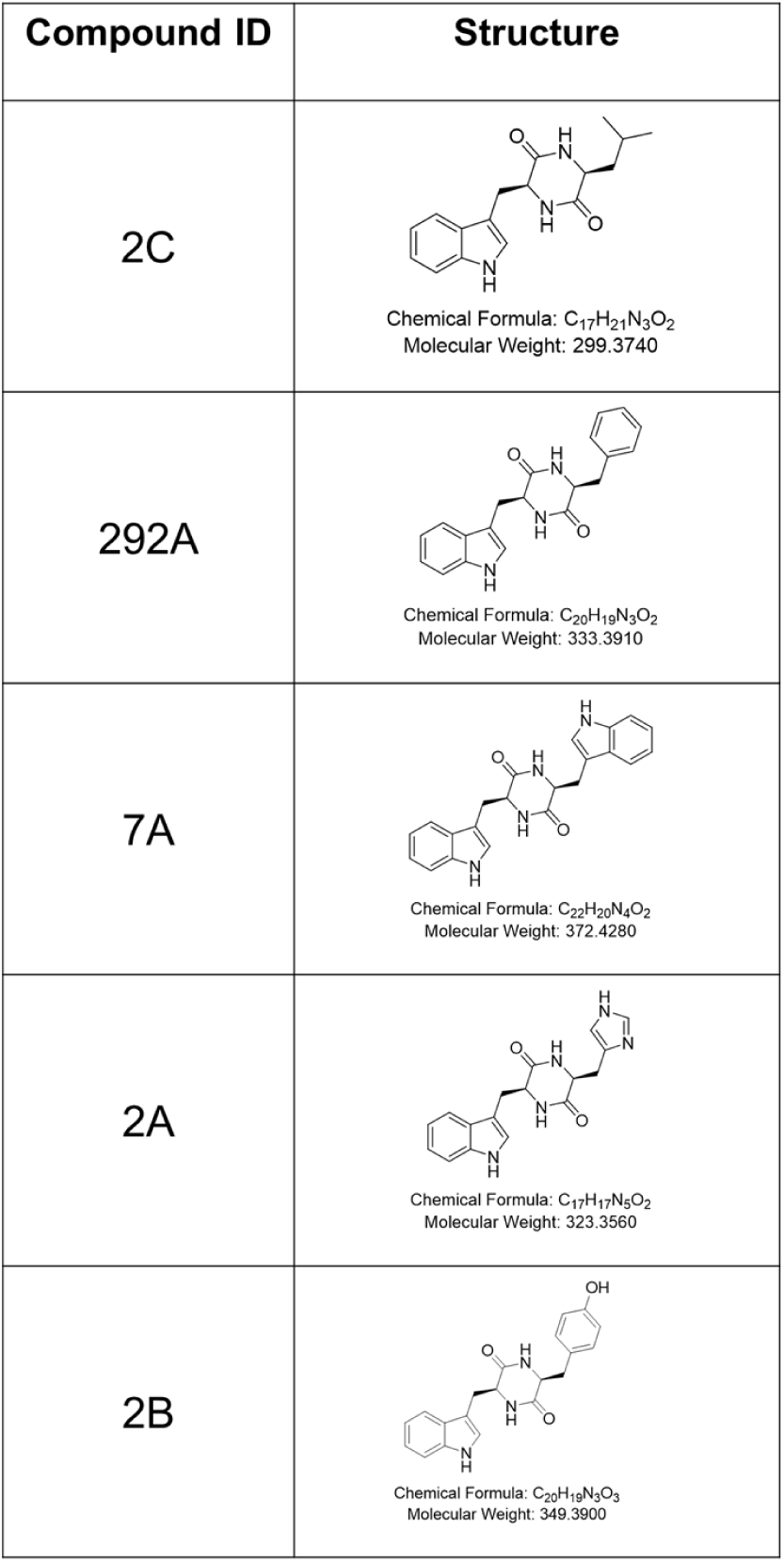
Summary of DKPs synthesized for culture supplementation.

Unlike soil extract and LPS, which showed their strongest effects at the 7 day timepoint, DKPs produced more significant changes at the 14 day timepoint. Moreover, their effects were concentration-dependent, with 10 μM treatments generally producing more significant responses than 100 nM (Fig 7 and Fig S8).

**Figure 7.**
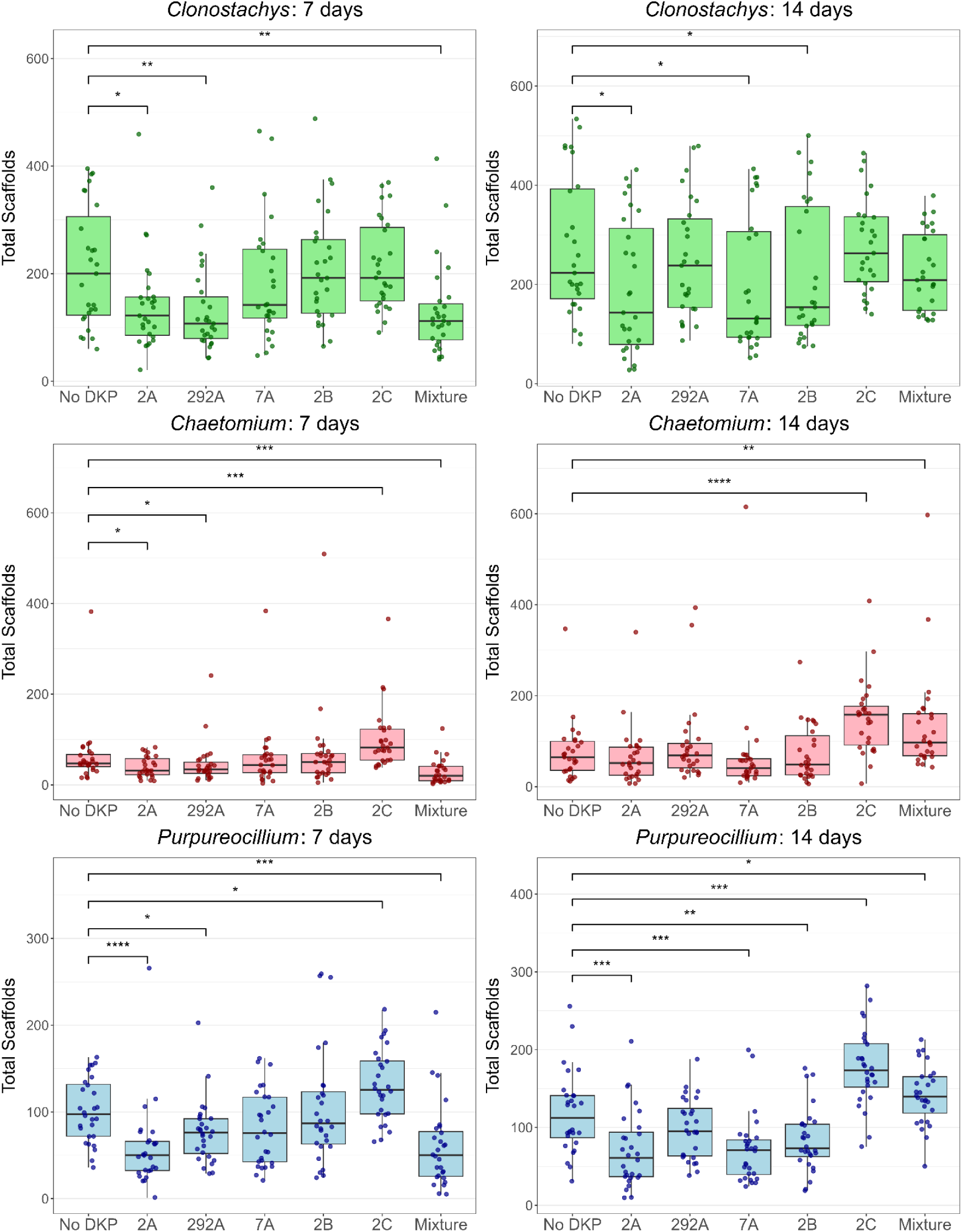
Summary of scaffold counts for 10 μM of DKP supplementation. Asterisks indicate statistical significance determined via Wilcoxon rank-sum test. * p < 0.05, ** p < 0.01, *** p < 0.001, **** p < 0.0001.

**Figure 8.**
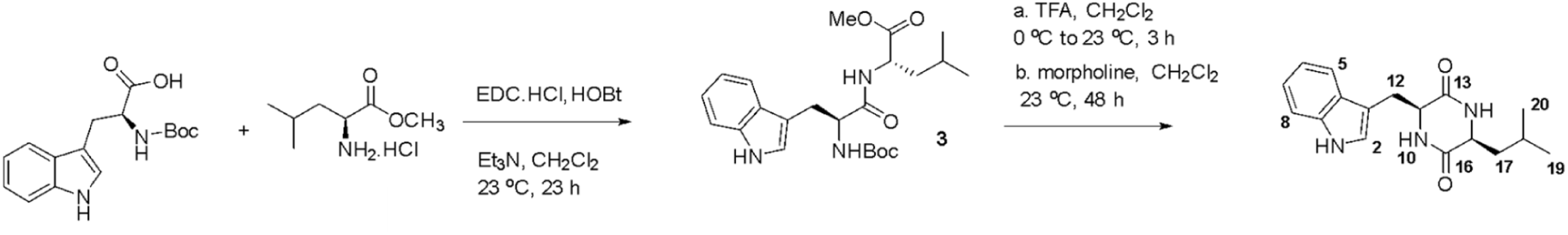
**Synthesis overview of DKP 2C.**

The impact of DKP supplementation on scaffold diversity varied depending on the compound and concentration. DKPs 2A and 2C tended toward exerting the most pronounced effects. At the 7-day time point, DKP 2A significantly decreased scaffold diversity, mirroring the suppressive effect observed with LPS. However, by 14 days, DKP 2A led to a significant increase in scaffold output, suggesting a time-dependent induction of biosynthetic activity. DKP 2C was the most effective stimulator overall, producing substantial increases in scaffold diversity at the 14-day time point.

Genera-specific responses were also observed. *Purpureocillium* cultures were highly sensitive to DKP supplementation, showing marked shifts in scaffold diversity for more DKPs, while *Clonostachys* exhibited the least amount of significant changes under any DKP condition.

Together, these results suggest that DKPs act as longer-term modulators of fungal secondary metabolism, capable of both suppressing and enhancing scaffold diversity depending on timing, compound identity, and concentration. Unlike environmental extracts such as soil or LPS, DKPs may offer a more controllable and tunable means of manipulating fungal metabolomes in crude extract library development.

## Discussion

Our analysis of fungal crude extract library design reveals that some rational strategies may not deliver the expected gains in chemical diversity. Across both geographic and phylogenetic selection strategies, we found no significant improvements over random sampling. Additionally, we conducted a similar analysis using morphological diversification as the basis for library design, focusing on diversification of color in different parts of fungal anatomy. However we observed similar outcomes, where it did not outperform random selection (Fig. S11)

This lack of improvement by rational selection compared to random selection is notable because it challenges a potentially intuitive assumption that maximizing phylogenetic distance or regional spread will maximize chemical space coverage. Much of the literature emphasizes rational selection approaches, but this is often influenced by positive-results bias, where only successful cases are reported. In practice, our results suggest that random selection is at least as effective as geographic or phylogenetic rational schemes, while also requiring less effort and prior information from researchers. This result offers exciting potential as it suggests that robust regionally-focused specimen collection efforts might be more effective than site-hopping across geographically distinctive habitats.

In contrast, culture conditions did influence scaffold diversity, though the fungi were not as broadly sensitive. Among the different conditions tested, DKP supplementation, particularly DKP 2C at 14 days, was the most effective, increasing scaffold diversity by 49%. Soil extract supplementation at 7 days was the second most effective, providing an early but transient boost in scaffold output (Table 3). LPS addition, meanwhile, yielded more nuanced effects, suppressing rather than stimulating scaffold production. These results highlight that environmental cues can shape metabolite profiles, but their effects are often genus-specific, time-dependent, or concentration-dependent, rather than universal. It is possible that supplementation, rather than stimulating unique metabolite production, may inhibit some biosynthetic genes. Alternatively, their additions may cause a stress response in the fungi, resulting in downregulation of total metabolite production.

**Table 3.** Average percentage change amongst scaffold counts for all fungal isolate culturing conditions. Standard error of the percent change is included. For a summary showing differences between each fungal genera, see Table S4.

|  | 7 day | 14 day |
| --- | --- | --- |
| Soil extract | 32% ± 12% | 3% ± 11% |
| Soil extract + sucrose | -25% ± 8% | -11% ± 11% |
| LPS -<br>1.8 ng/ml | -39% ± 7% | 6% ± 14% |
| DKP -<br>2A (10 μM) | -38% ± 12% | -28% ± 14% |
| DKP -<br>292A (10 μM) | -30% ± 13% | -2% ± 17% |
| DKP -<br>7A (10 μM) | -13% ± 17% | -27% ± 17% |
| DKP -<br>2B (10 μM) | 4% ± 22% | -18% ± 15% |
| DKP -<br>2C (10 μM) | 31% ± 21% | 49% ± 22% |
| DKP -<br>Mixture (10 μM) | -44% ± 11% | 26% ± 21% |

We hypothesized that chemical diversity may be strongly associated with the amount of fungal biomass produced under each culture condition. Although biomass was not directly measured in these experiments, raw MS signal intensity from fungal-derived features was used as a proxy for biomass. Correlations were constructed between total feature count and total raw MS signal, and these relationships were consistently strong across timepoints and genera within a given fungal genus (Figs S12–S15). These results indicate that total biomass can be used to approximate chemical diversity in a fungal culture.

Taken together, our findings suggest a path forward for smarter fungal library design. Selection strategies based on geography or phylogeny appear inefficient for expanding chemical space, whereas cultivation strategies that mimic environmental signals can provide measurable gains, albeit in a targeted and context-dependent manner. By systematically comparing both approaches with high-resolution tandem mass spectrometry, we show that intuitive strategies do not always align with chemical outcomes. Future efforts should therefore prioritize cultivation-based interventions, such as those covered by our supplementation data and other studies that also follow OSMAC principles, rather than investing heavily in rational selection schemes that may not outperform random sampling strategies.

## Acknowledgements

Research reported in this publication was supported by the National Institute Of General Medical Sciences of the National Institutes of Health under Award Number R01GM145649. The content is solely the responsibility of the authors and does not necessarily represent the official views of the National Institutes of Health. Authors wish to thank C. Brown, C. Miller and J. Rangel for technical assistance with fungal cultures, morphology analyses and PCR.

## Methods

### Fungal collection

All fungal extracts were isolated from soil samples sent through the University of Oklahoma Citizen Science Program ^28^. For all extracts, the soil sample, zipcode, and time of collection were recorded. For the *Trichoderma*-isolates, extra details including the texture and color of fungal characteristics like hyphae, mycelium, pustules, etc, were recorded.

### Diketopiperazine synthesis methods

For more detailed synthesis and spectra-see Supplemental Data 1. Below is a brief summary.

All Diketopiperazines were synthesized following the protocol developed by Koshizuka et al ^29^. Briefly, tert-butoxycarbonyl)-L-tryptophan (1 equiv), 1-hydroxybenzotriazole hydrate (1.5 equiv), and a methyl ester hydrochloride whose amino acid residue matched the variation of the DKP (1.5 equiv) were dissolved in dichloromethane. Triethylamine (4.5 equiv) was added to the mixture. After 10 minutes, EDC.HCl (1.5 equiv) was added and the resulting mixture was stirred for 8 h before another portion of EDC.HCl (0.6 equiv) and triethylamine (2 equiv) were added. The reaction mixture was stirred for an additional 15 h.

The mixture was washed with an aqueous solution of HCl and saturated aqueous NaHCO_3_ solution. The organic layer was dried over anhydrous Na_2_SO_4_, filtered and was concentrated in vacuo to afford a yellow foam. The crude was purified using a silica column.

The resulting dipeptide was dissolved in dichloromethane and trifluoracetic acid was added at 0 °C. The volatiles were removed after 3 h and the residue was redissolved in dichloromethane. To the resulting mixture, morpholine was added and the solution was vigorously stirred for 48 h as the product was precipitated as a white solid. The product was filtered under reduced pressure and washed with DCM and deionized water to afford a solid. The product was further purified using preparative HPLC, C18 column using 80% ACN/H2O with 0.1% FA.

### Fungal culturing and extraction

These fungal isolates were cultured on a solid-state medium composed of Cheerios breakfast cereal supplemented with a 0.3% sucrose solution containing 0.005% chloramphenicol, as previously optimized ^30,31^. The samples from the broad phylogeny and *Trichoderma* runs were cultured for 3 weeks. The soil extract, lipopolysaccharide, and diketopiperazine additions were cultured for either 7 or 14 days, depending on the experiment groups.

For isolates with soil extract addition, 7 kg of soil was collected from a wooden area owned by the University of Oklahoma (GPS coordinates 35.18,-97.44) in August of 2023. Water was added 1:1 vol:vol and autoclaved twice to ensure sterility. The aqueous extract was then evaporated *in vacuo* and the resulting extract was added to the fungal cultures at 15 g/L when the cultures were started.

For the lipopolysaccharide experiments, lipopolysaccharide (Sigma L3129) was resuspended in Millipore water at 1000x concentration. Diketopiperazines were resuspended in DMSO at 1000x stock concentration. Both were added to the fungal cultures at the indicated concentrations when the cultures were started All fungal extracts were prepared using the protocol previously described ^15^. Briefly, fungal cultures in 16 × 100 mm borosilicate tubes underwent two successive water:ethyl acetate (1:1, v/v) partitions. The aqueous phase was discarded, and the organic phase was evaporated in vacuo. Residues were stored at −20 °C and later resuspended in DMSO for screening. For LC-MS/MS analysis, extracts were diluted 1:10 in methanol containing 2 μM sulfadimethoxine as an internal standard.

### Sequencing

For the sequence generation and phylogenetic analysis for *Trichoderma,* the internal transcribed spacer (ITS) region of the isolates was PCR amplified and sequenced as previously reported ^32^ (Genbank accession numbers PV347956 - PV348905). ITS sequences from three fungi differing at the phylogenetic class level from *Trichoderma* were included to function as an outgroup (Genbank MT723856.1, KU179108.1, OP965332.1). Seven published *Trichoderma* ITS sequences were included to show phylogenetic relationships within the genus (Genbank NR_145035.1, AF487658.1, FJ904855.1, AY380903.1, AY605713.1, NR_138448.1, AY380905.1).

For each isolate, a ∼600 bp portion of the translation elongation factor 1-alpha gene (tef) was amplified and sequenced (Genbank accession numbers PV344763-PV345453). Primers and parameters used were previously published ^32^. Briefly, EF1-1018F and EF1-1620R were used in a touchdown PCR with annealing temperatures from 66°C to 56°C followed by 36 cycles at 56°C. tef sequences from two *Dothideomycetes* fungi were used to root the tree (Genbank PP764432.1, MW25816.1). Seven *Trichoderma* tef sequences were included to show phylogenetic relationships within the genus (Genbank OQ116869.1, DQ835481.1, OR333860.1, OM972899.1, OQ988162.1, OP970252.1, OP970249.1).

Raw sequencing data were converted to fastq files in Biopython ^33^ and contigs were assembled in mothur ^34^. Resulting sequences were aligned in ClustalW ^35^ and manually trimmed. Distance matrices were made in MEGA X ^36^, with the maximum composite likelihood and 500 bootstraps. Phylogenetic analyses were conducted in R using the phangorn, ape, and SeqinR packages ^37–39^. These generated a maximum likelihood tree with 500 bootstrap replicates for both genes. Trees were visualized via Evolview ^40^.

### Mass spectrometry analysis

The multi-genera collection data presented is a re-analysis of a collection previously described and analyzed ^17^.

After resuspension in 1:10 in methanol containing 2 μM sulfadimethoxine as an internal standard., samples were separated using a Kinetex 1.7 µm C18 50 x 2.1 mm LC column with a C18 guard cartridge (Phenomenex) on a Thermo Scientific Vanquish ultra-high-performance liquid chromatography (UHPLC) instrument using only MS-grade solvents (Fisher Optima). The following gradients were used (Tables 4-5)

**Table 4.** Liquid chromatography gradient summary, used for multi-genera data collection.

| Time | Mobile phase |
| --- | --- |
| 0-1 min | 5% B |
| 1-8 min | Increase to 100% B |
| 10-10.5 min | 100% B |
| 10.5-12.5 | Decrease to 5% B |
| A: Water + 0.1% v/v formic acid<br>B: Acetonitrile + 0.1% v/v formic acid<br>Flow rate of 0.5 ml/min used throughout gradient |  |

**Table 5.** Liquid chromatography gradient summary, used for *Trichoderma*, Soils, LPS, and diketopiperazine data acquisition.

| Time | Mobile phase |
| --- | --- |
| 0-1 min | 5% B |
| 1-9 min | Increase to 100% B |
| 9-10.5 min | 100% B |
| 10.5-12.5 | Decrease to 5% B |
| A: Water + 0.1% v/v formic acid<br>B: Acetonitrile + 0.1% v/v formic acid<br>Flow rate of 0.5 ml/min used throughout gradient |  |

A Thermo Scientific Q-Exactive Plus was used for MS/MS analysis on the multi-genera collection, *Trichoderma*, soil extract, and LPS, and DKP analysis runs.

Only positive ionization data was collected for all extracts. The parameters used for MS collection are listed in Tables S5 and S6. Prior to analysis, the Q Exactive Plus was calibrated using Pierce LTQ Velos ESI positive ion calibration solution (ThermoFisher), and the Exploris 240 was calibrated using Flexmix (ThermoFisher).

For each run, the sample injection order was randomized within each plate, and the order of plate analysis was randomized. Every 12 samples, a solvent blank was injected and a pooled extract was injected to monitor instrument stability and potential batch effects. A standard mix of sulfadimethoxine, sulfachloropyridazine, sulfamethazine, sulfamethizole, amitriptyline, and coumarin-314 was injected every 100 samples to monitor retention time shifts.

### Data processing

Files were converted to mzML format using MSConvert ^41^. The feature table for all MS data was done using MZmine 2.53 using the parameters listed in Table S7.

Oversplit, broad background peaks were removed. Features appearing in the blank injections or cheerio media were removed using a 5-fold blank removal cutoff. For the environmental-mimicking cultures (soil extract, LPS, DKPs), features associated with the additions were removed under a 10-fold removal cutoff. All feature tables were normalized using the Total Ion Current (TIC) method. All codes used for data cleaning were run in Rstudio.

Some MS runs used slightly altered methods, particularly with regards to the *m/z* range used (Table S6). To ensure consistency in the data, all MS runs *m/z* range was truncated to only include *m/z* values between 120 and 1050.

Chemical families and metabolite annotations were found using Feature-Based Molecular Networking using GNPS1 ^14^. The parameters and spectral libraries used for analysis are listed in Tables S8 and S9.

For the phylogeny selection analysis, the selections were done using the distance matrices generated by the ITS and tef sequencing (See Methods: Sequencing). For the accumulation curves, each rational selection began with a random isolate selection. Using the distance matrix, the isolate with the highest distance from the starting isolate was added to the accumulation curve. This process repeats until all isolates have been added to the accumulation curve, always excluding isolates which have already been chosen from the curve. Twenty-five iterations are done for the curve.

Random Forest based analysis was performed on peak area data derived from *Chaetomium*, *Purpureocillium*, and *Clonostachys* samples collected at 7-day and 14-day time points under treatment with 1.8 ng of LPS, with comparison to the matched timepoint in the absence of LPS treatment. An additional comparison was performed between 7-day and 14-day timepoints in the absence of LPS, to identify effects of culturing time. Initial feature classification was performed in R using the Boruta Random Forest algorithm (doTrace = 2, maxRuns = 100, set.seed(123)). To further evaluate the predictive relevance of the selected features, model validation was performed in Python using the XGBoost library. The dataset was split into training and testing subsets using a stratified 70/30 split (random_state = 123) to preserve class balance. Model performance was assessed using repeated stratified K-fold cross-validation, and all models achieved a mean accuracy >0.80. Following validation, features were stratified by taxonomic group and time point to identify metabolites uniquely affected in each condition. Scripts were developed in Jupyter Notebook using R version 4.4.1 and Python 3.12.4. These unique features were subsequently merged with chemical family annotations identified through GNPS1.

## Data availability

The LC-MS data is available under the following massive repositories (Table 6).

**Table 6.** MS data availability and analysis links.

| Run | Massive Dataset ID | GNPS Feature-Based Molecular Networking Link |
| --- | --- | --- |
| Multi-genera collection | MSV000091950 | https://gnps.ucsd.edu/ProteoSAFe/status.jsp?task=6d82b4b5607d429cad0afa5061e67235 |
| *Trichoderma* | MSV000092854 | https://gnps.ucsd.edu/ProteoSAFe/status.jsp?task=7cc8217463cb49ae8926b3b418cf1072 |
| Soils/LPS | MSV000093972 | https://gnps.ucsd.edu/ProteoSAFe/status.jsp?task=fd416e4cf20e4c2d9426beee7a835cd5 |
| Diketopiperazines | MSV000094173 | https://gnps.ucsd.edu/ProteoSAFe/status.jsp?task=e1fe0f695dba4cdfb24c132d7d52aca8 |

The Molecular Networking links are also listed. Genbank codes are available for *Trichoderma* ITS (PV347956 - PV348905), *Trichoderma* tef (PV344763-PV345453), and environmental mimicking isolates (PX249573-PX249656). XGBoost code can be accessed from: https://github.com/demetrius-tillery/McCall-Lab/tree/main/Natural%20Products%20Work

## Supplemental Data

**Supplemental Data 1**. Detailed Synthesis and spectra for the 5 diketopiperazines used for culture supplementations

## Figures

**Figure S1.**
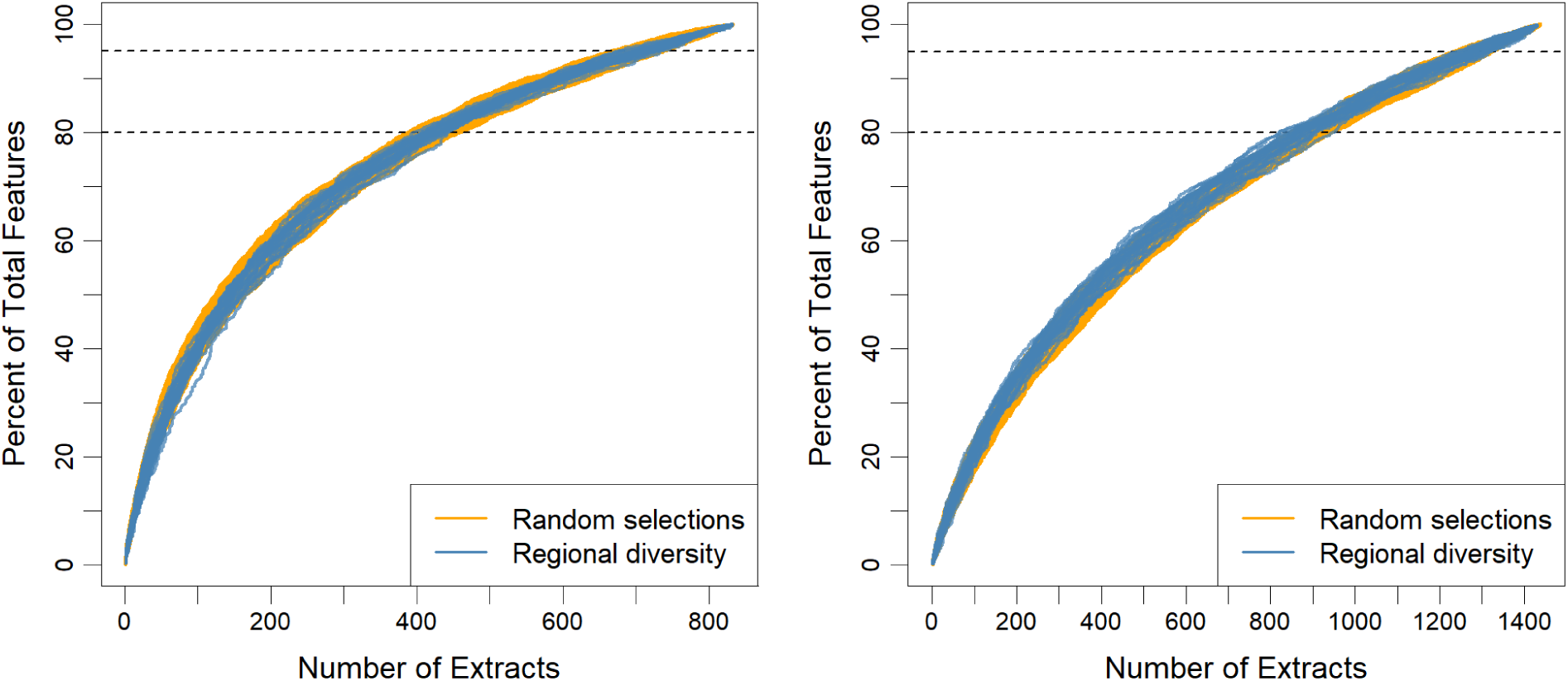
Regional diversity prioritization does not outperform random selection - feature level. Feature level analysis of rational selection based on regional diversity. This analysis mirrors that shown in Figure 1 but is performed at the feature level rather than the scaffold level.

**Figure S2.**
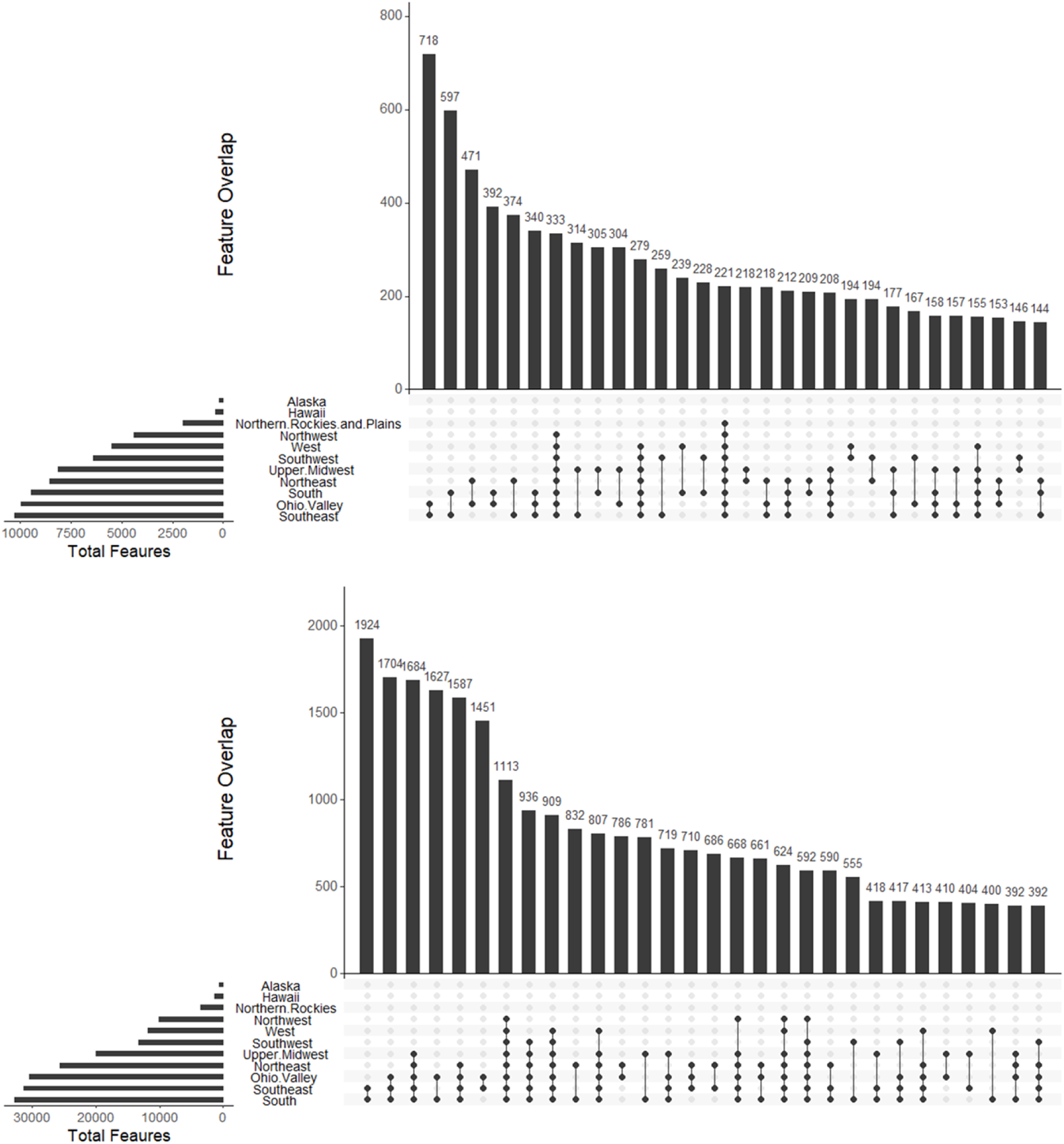
Broad overlap of shared features across regions. UpSet plots showing the distribution of features shared among geographic regions. Only the top 30 intersection sets are displayed, and features detected in a single region only are excluded. The top panel shows results for *Trichoderma*, while the bottom panel shows the multi-genera dataset. This analysis mirrors that shown in Figure 2 but is performed at the feature level rather than the scaffold level.

**Figure S3.**
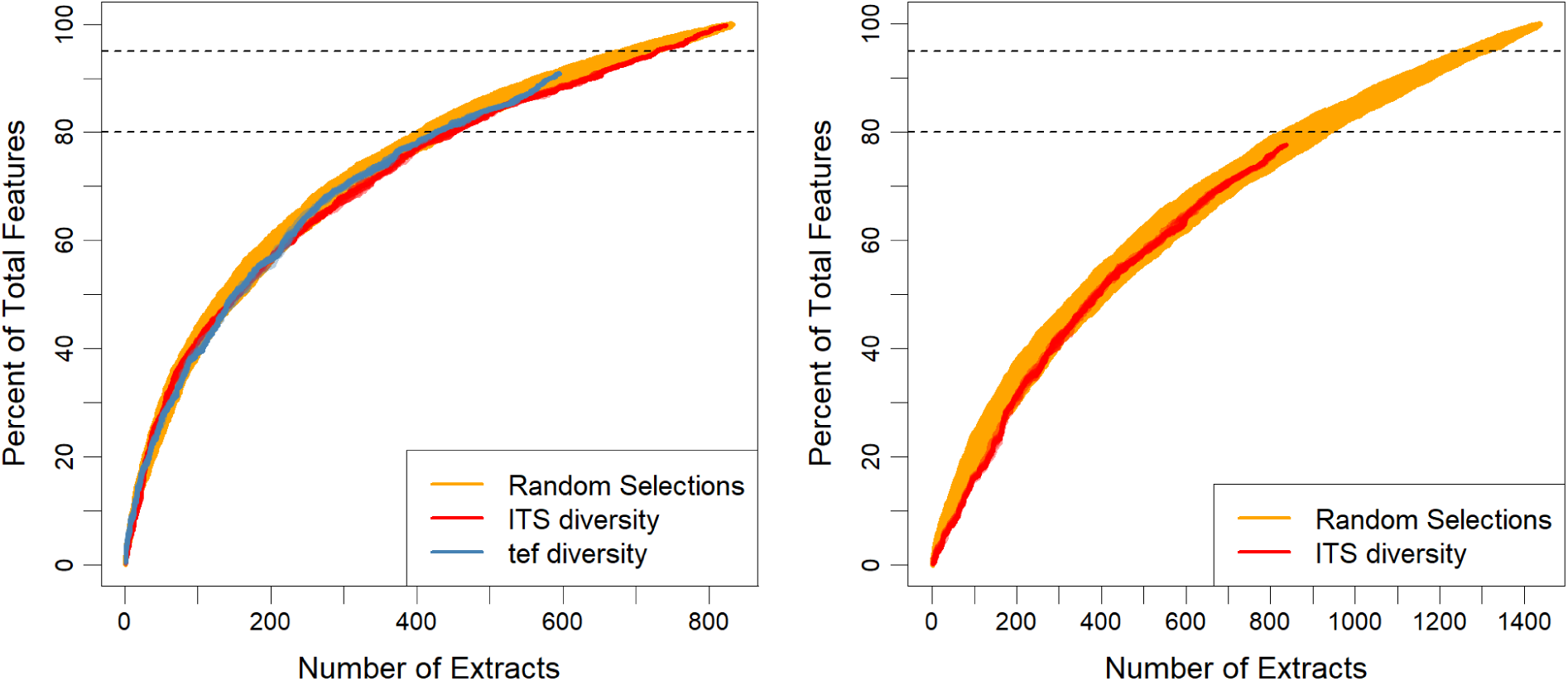
Phylogeny-based selection does not outperform random selection - feature level. Each rational phylogeny-based selection started with a random isolate, and subsequent isolates were selected to maximize phylogenetic diversity. The results for the *Trichoderma* collection are shown on the left, and the multi-genera collection is shown on the right. For both datasets, 25 iterations were used. This analysis mirrors that shown in Figure 3 but is performed at the feature level rather than the scaffold level.

**Figure S4.**
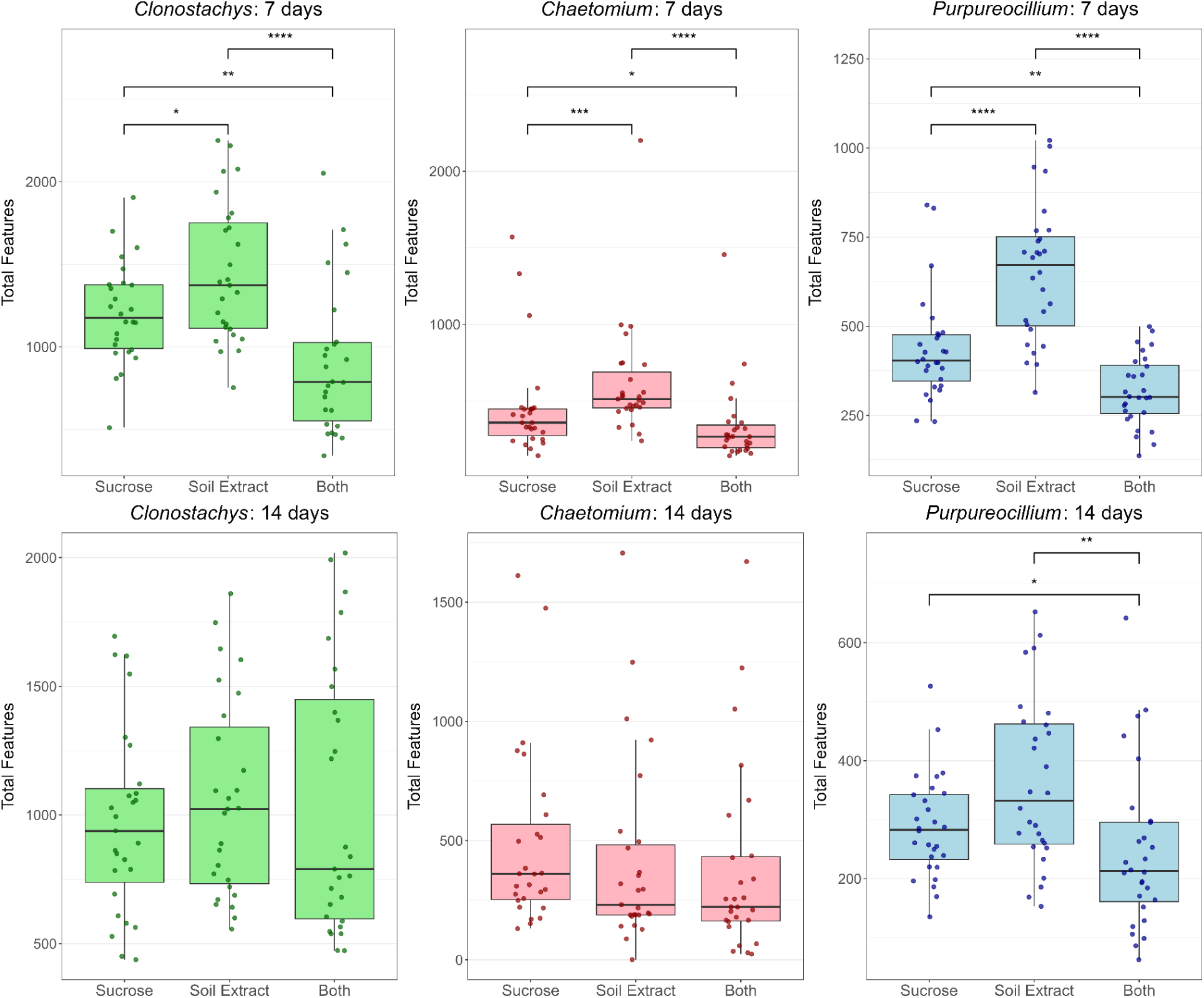
Summaries of total features per isolate, separated by genera and timepoint - feature level. Asterisks indicate statistical significance determined via Wilcoxon rank-sum test. * p < 0.05, ** p < 0.01, *** p < 0.001, **** p < 0.0001. This analysis mirrors that shown in Figure 4 but is performed at the feature level rather than the scaffold level.

**Figure S5.**
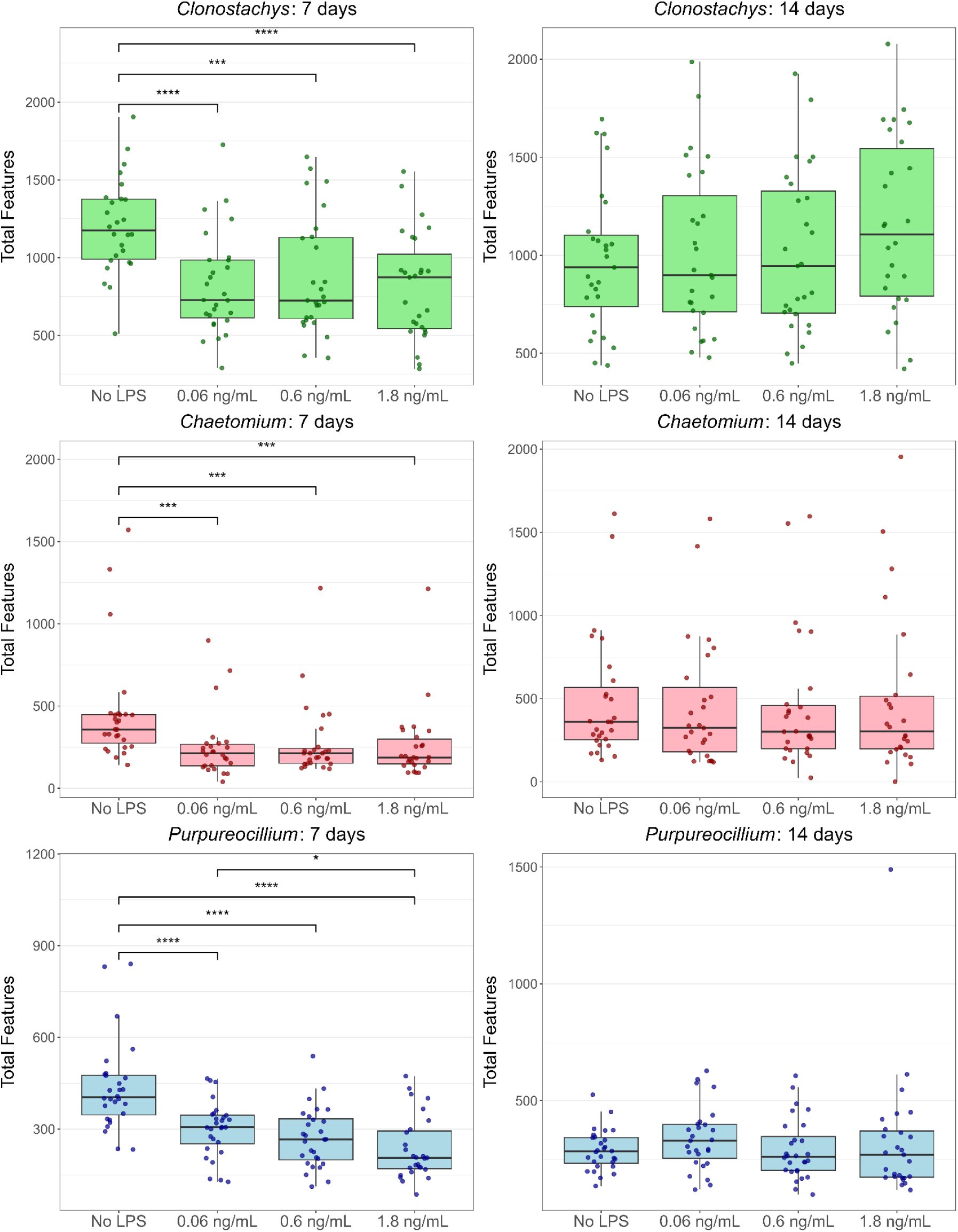
Boxplots showing feature count per isolate for each genera, timepoint, and LPS experimental group. Asterisks indicate statistical significance determined via Wilcoxon rank-sum test. * p < 0.05, ** p < 0.01, *** p < 0.001, **** p < 0.0001. This analysis mirrors that shown in Figure 5 but is performed at the feature level rather than the scaffold level.

**Figure S6.**
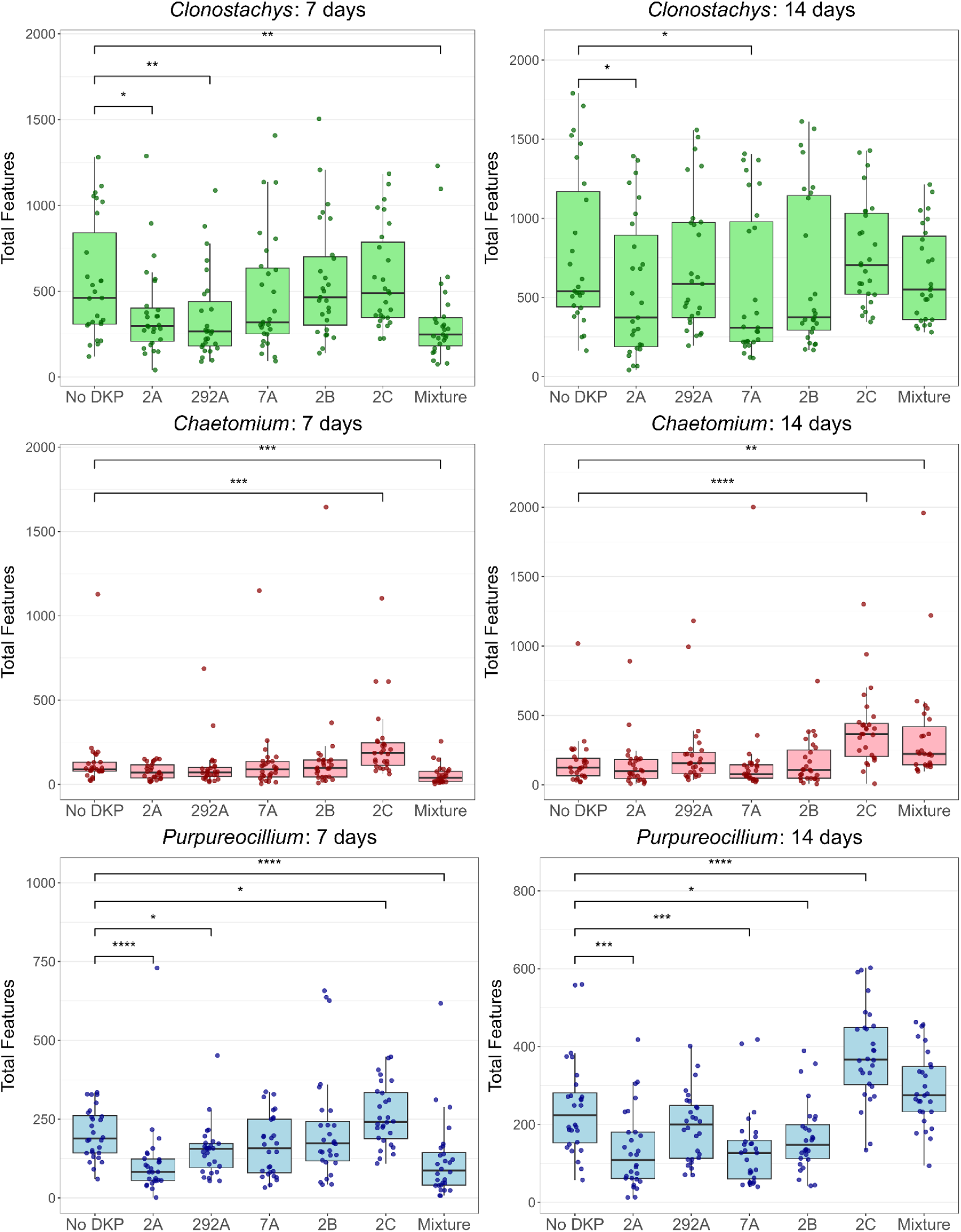
Summary of feature counts for 10 μM of DKP supplementation. Asterisks indicate statistical significance determined via Wilcoxon rank-sum test. * p < 0.05, ** p < 0.01, *** p < 0.001, **** p < 0.0001. This analysis mirrors that shown in Figure S8 but is performed at the feature level rather than the scaffold level.

**Figure S7.**
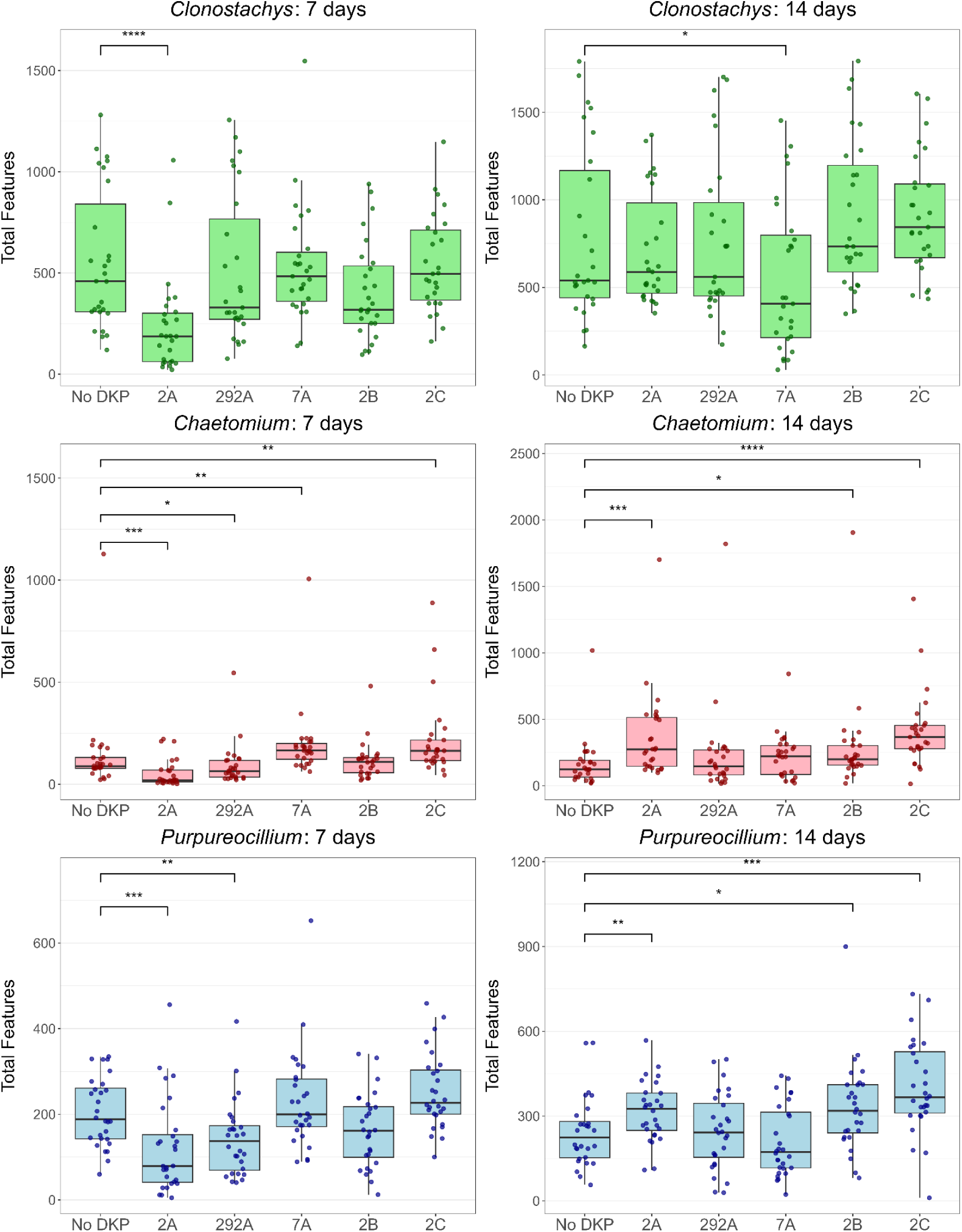
Summary of feature counts for 100 nM of DKP supplementation. Asterisks indicate statistical significance determined via Wilcoxon rank-sum test. * p < 0.05, ** p < 0.01, *** p < 0.001, **** p < 0.0001. This analysis mirrors that shown in Figure 7 but is performed at the feature level rather than the scaffold level.

**Figure S8.**
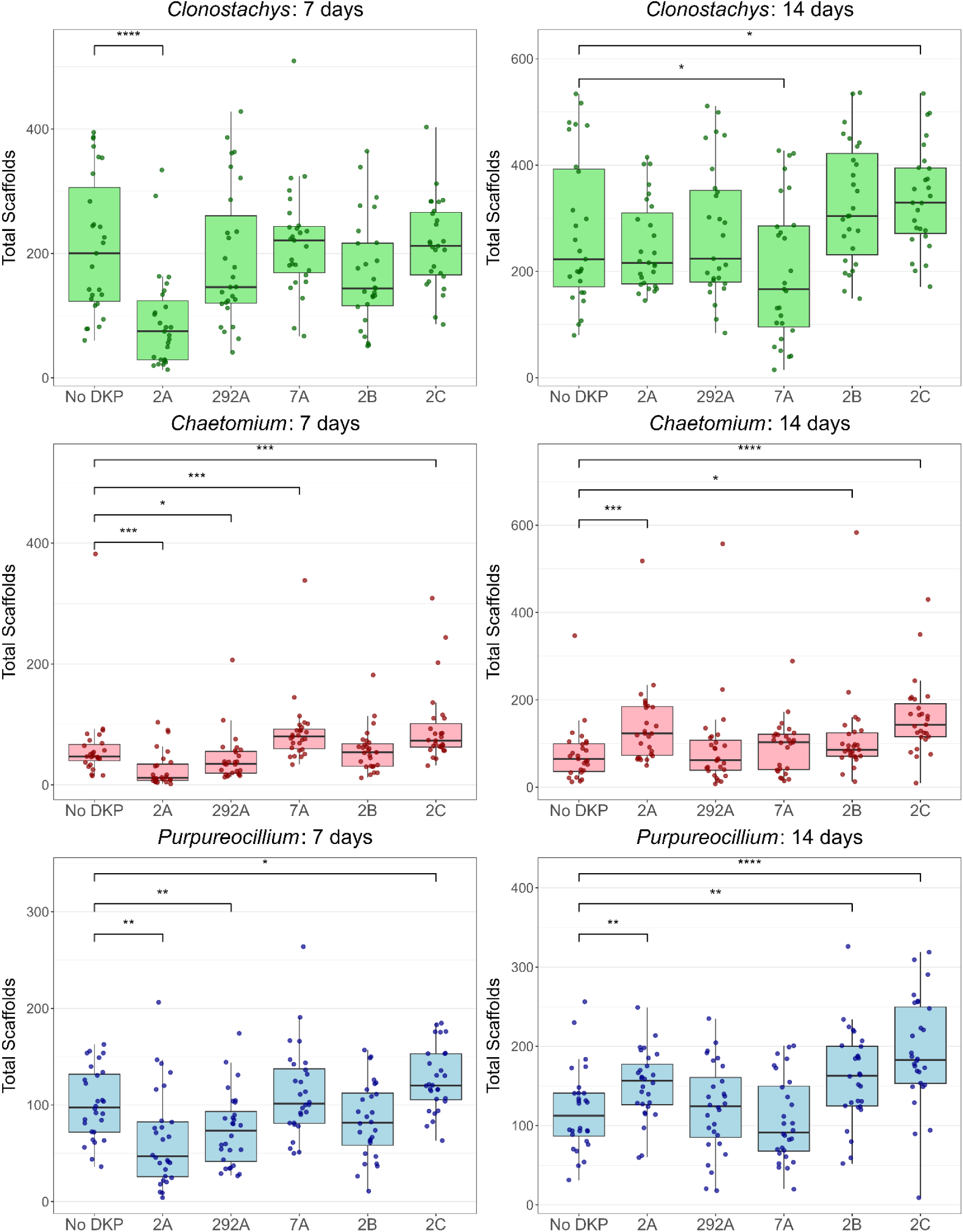
Summary of scaffold counts for 100 nM of DKP supplementation. Asterisks indicate statistical significance determined via Wilcoxon rank-sum test. * p < 0.05, ** p < 0.01, *** p < 0.001, **** p < 0.0001.

**Figure S9.**
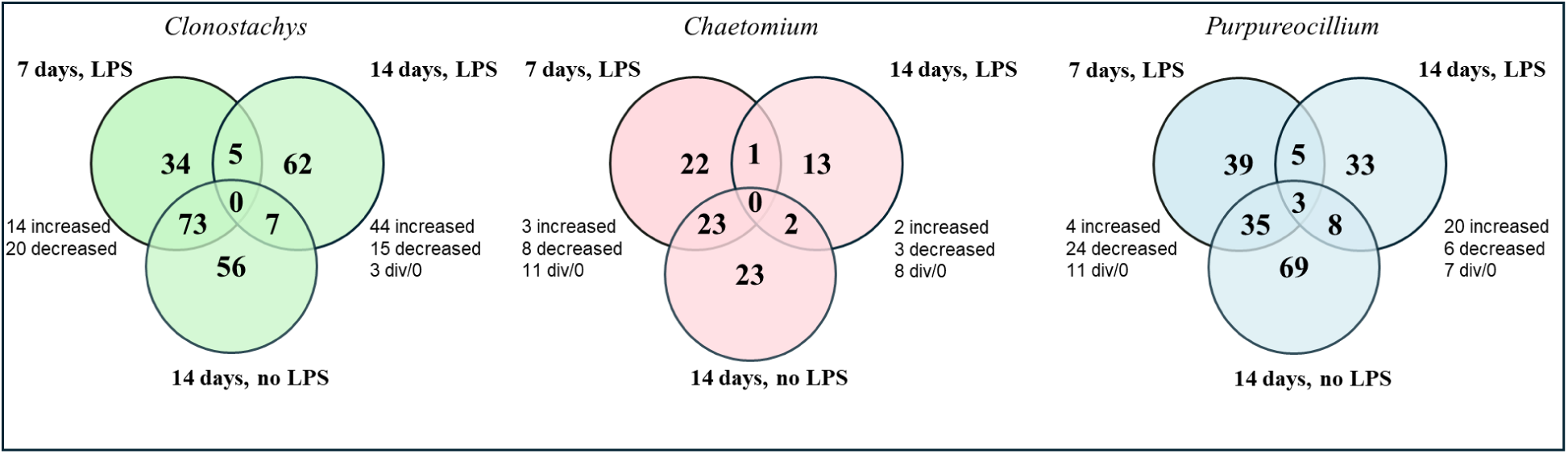
**Significant features impacted by culture time and LPS treatment**. Each venn diagram represents a comparison of significant features of the genera *Clonostachys*, *Chaetomium*, or *Purpureocillium*. Through Boruta random forest analysis, all peak area data for a specific genus at 7 days without LPS treatment was compared to 7 days with 1.8 ng of LPS, 14 days with 1.8 ng LPS, or 14 days without LPS treatment.

**Figure S10.**
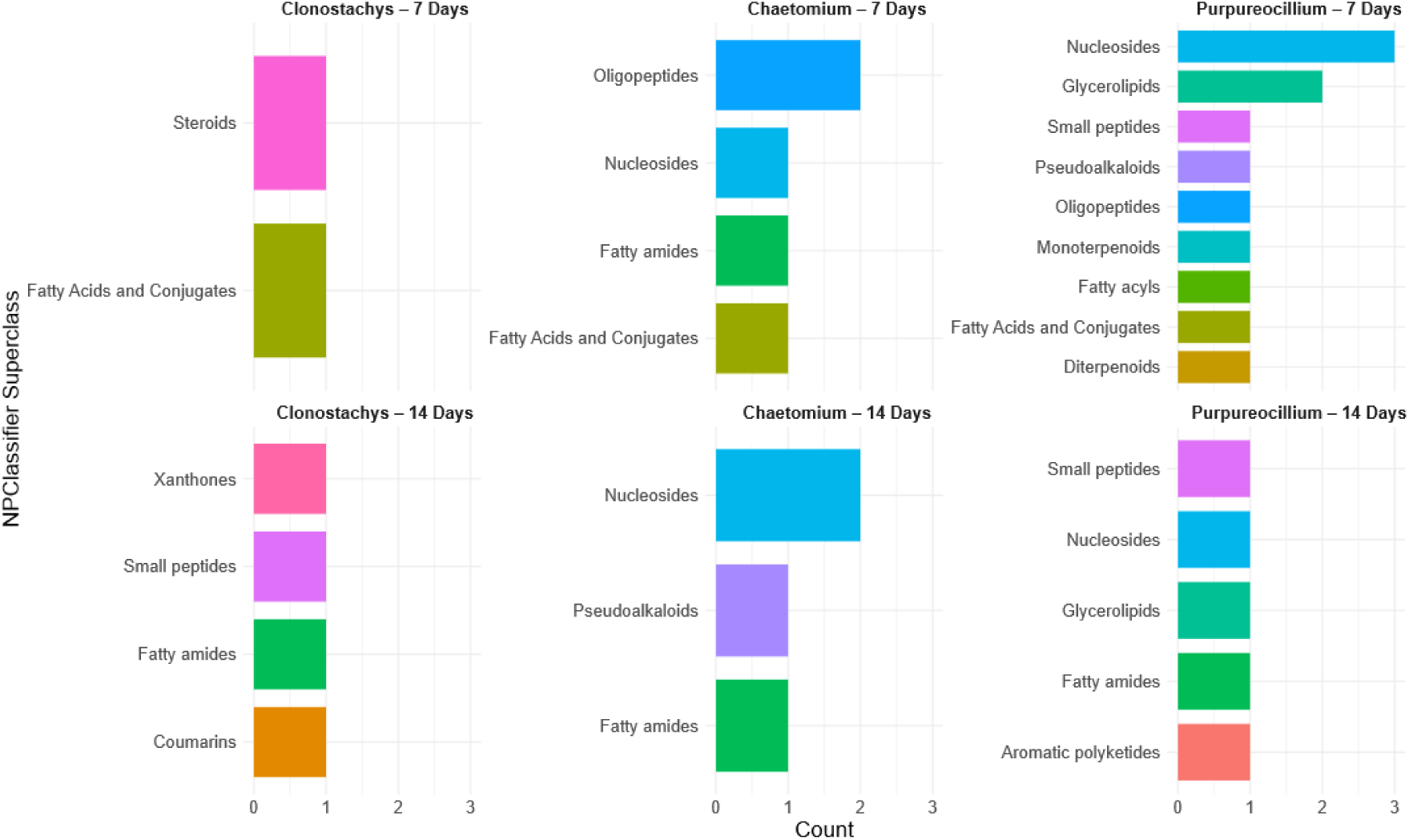
**Comparisons of annotated chemical superclasses that experienced significant changes**.For each genera (*Clonostachys*, *Chaetomium*, and *Purpureocillum)* and timepoint (7 and 14 days). Annotated chemical superclasses that were significantly altered are shown. These superclasses represent metabolite groups that were differentially regulated in response to 1.8 ng/mL LPS supplementation compared to the unsupplemented control. Analysis determined via Boruta random forest.

**Figure S11.**
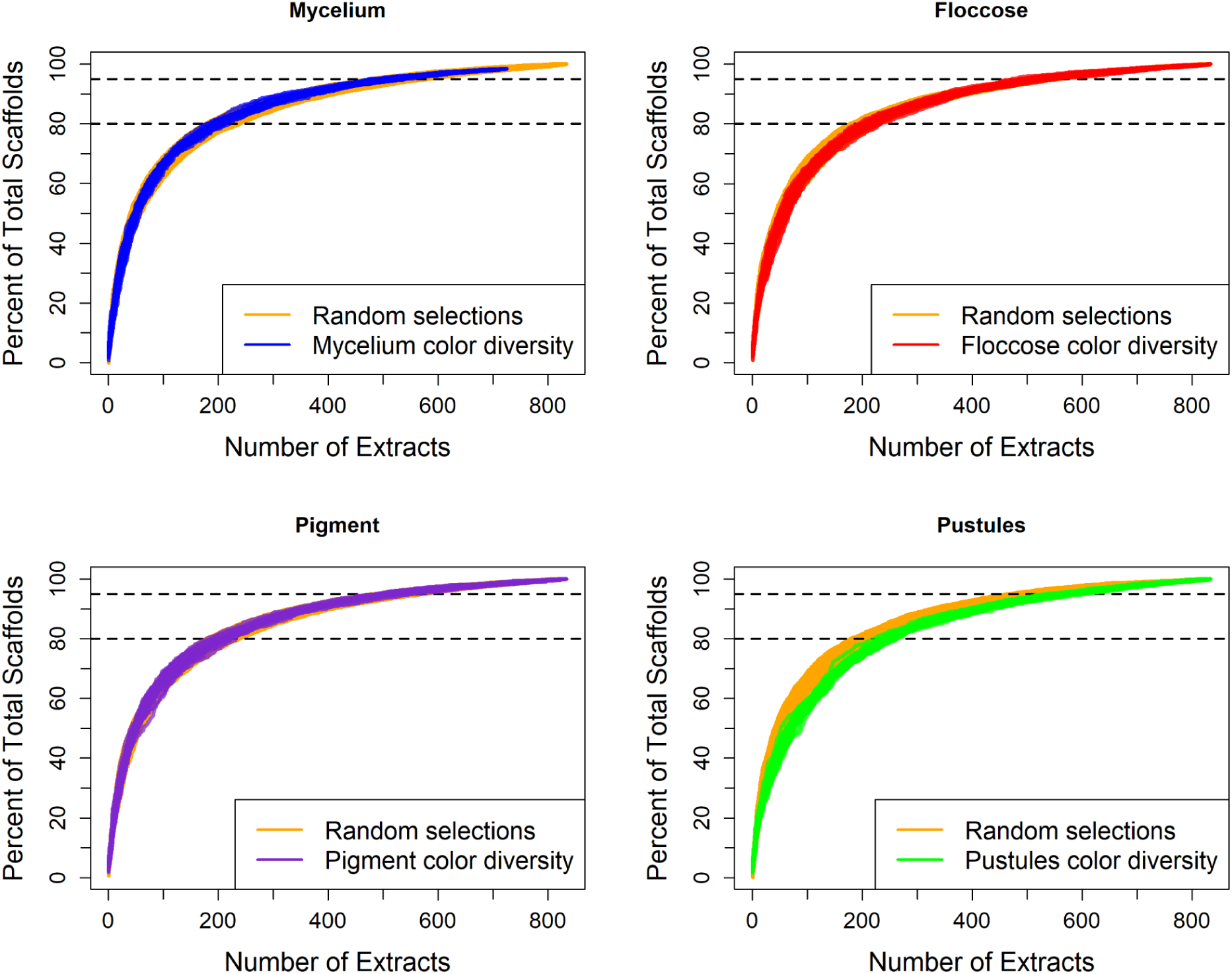
Morphological diversity prioritization does not outperform random selection. In the *Trichoderma* collection, each isolate was annotated for the color of multiple culture features, including mycelium, floccose growth, pigment, and pustules. Accumulation curves for morphology-based rational selection were generated by first selecting a random isolate, then iteratively adding isolates with colors underrepresented in the existing selection until all isolates were exhausted. Each analysis was repeated for 25 iterations.

**Figure S12.**
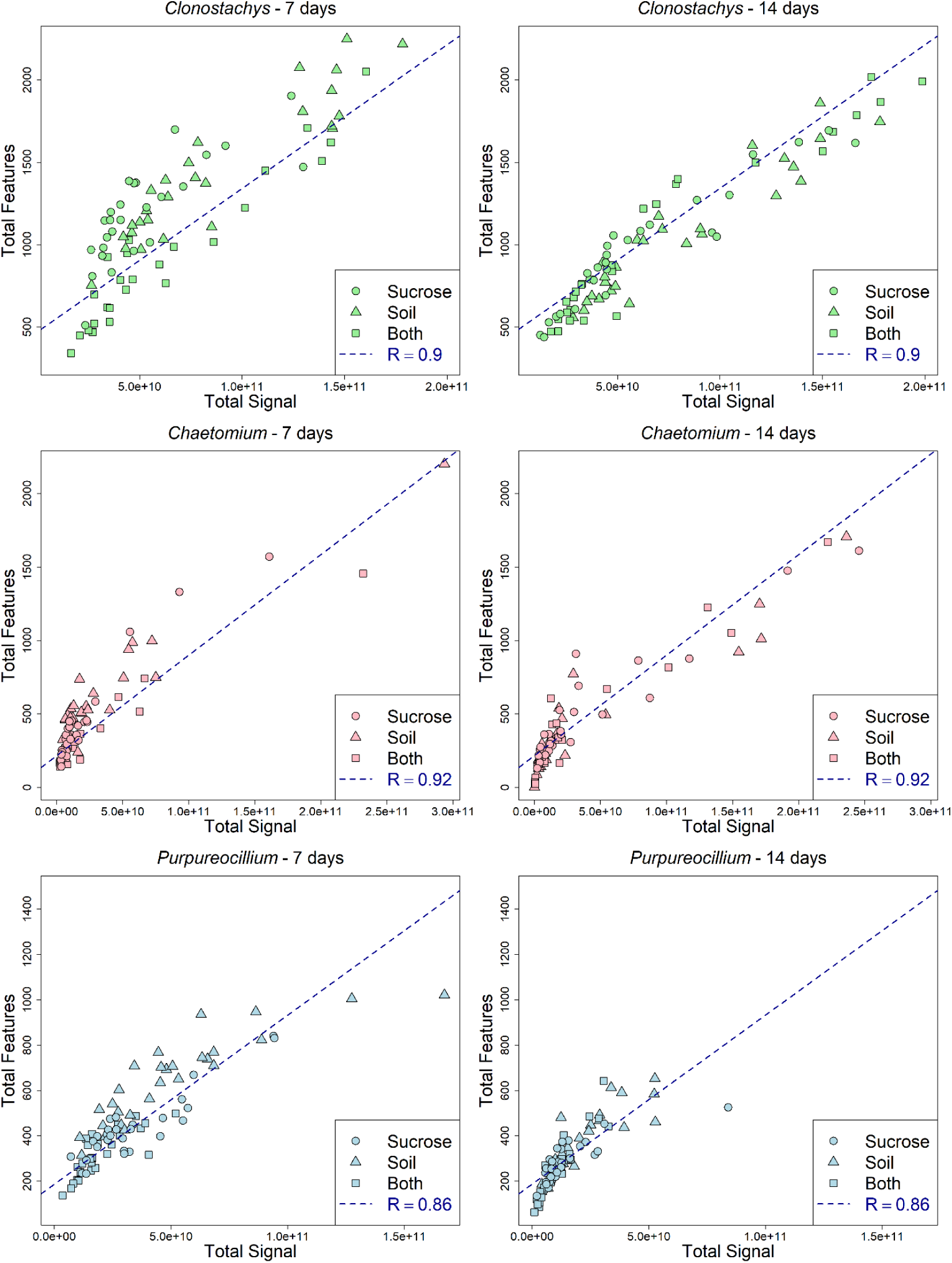
Total raw MS signal strongly correlates to chemical diversity - soil extract analysis. For each timepoint and genus, a correlation between raw MS signal (after 5-fold blank and media feature removal)(See Data Processing) and total feature count was built. Raw MS signal was used as a proxy for biomass for this analysis.

**Figure S13.**
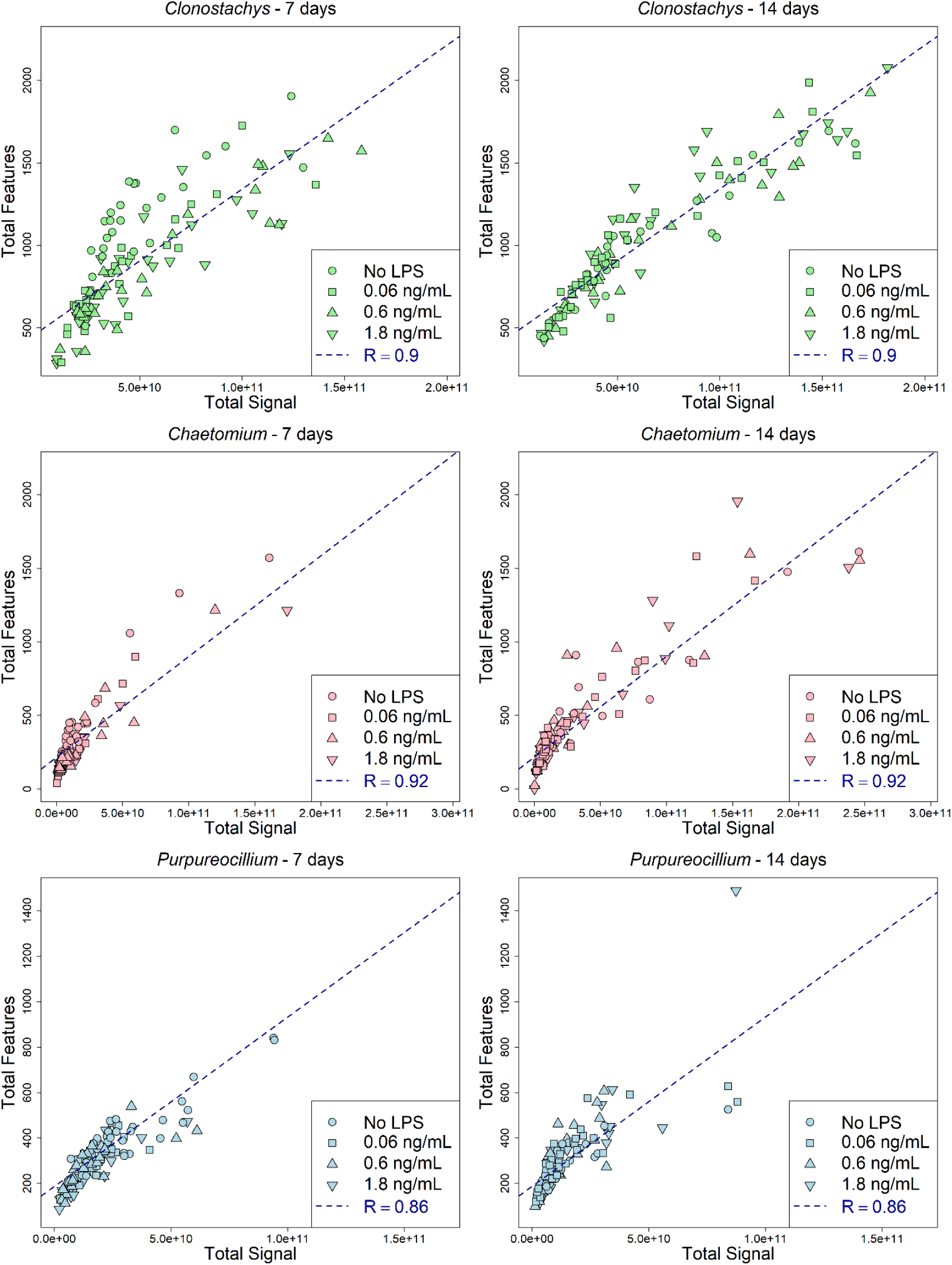
Total raw MS signal strongly correlates to chemical diversity - LPS supplementation analysis. For each timepoint and genus, a correlation between raw MS signal (after 5-fold blank and media feature removal)(See Data Processing) and total feature count was built. Raw MS signal was used as a proxy for biomass for this analysis.

**Figure S14.**
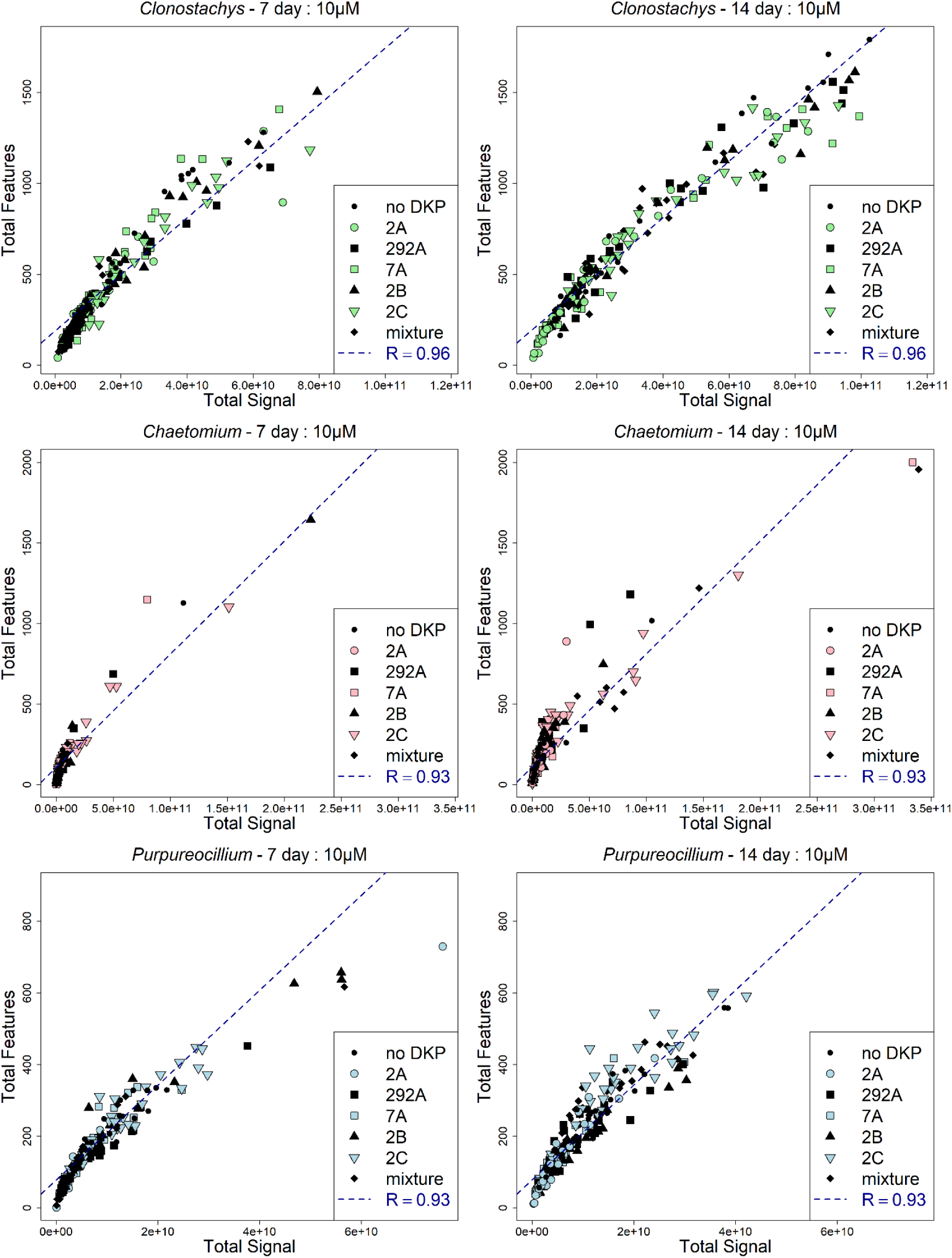
Total raw MS signal strongly correlates to chemical diversity - DKP 10μM supplementation. For each timepoint and genus, a correlation between raw MS signal (after 5-fold blank and media feature removal)(See Data Processing) and total feature count was built. Raw MS signal was used as a proxy for biomass for this analysis.

**Figure S15.**
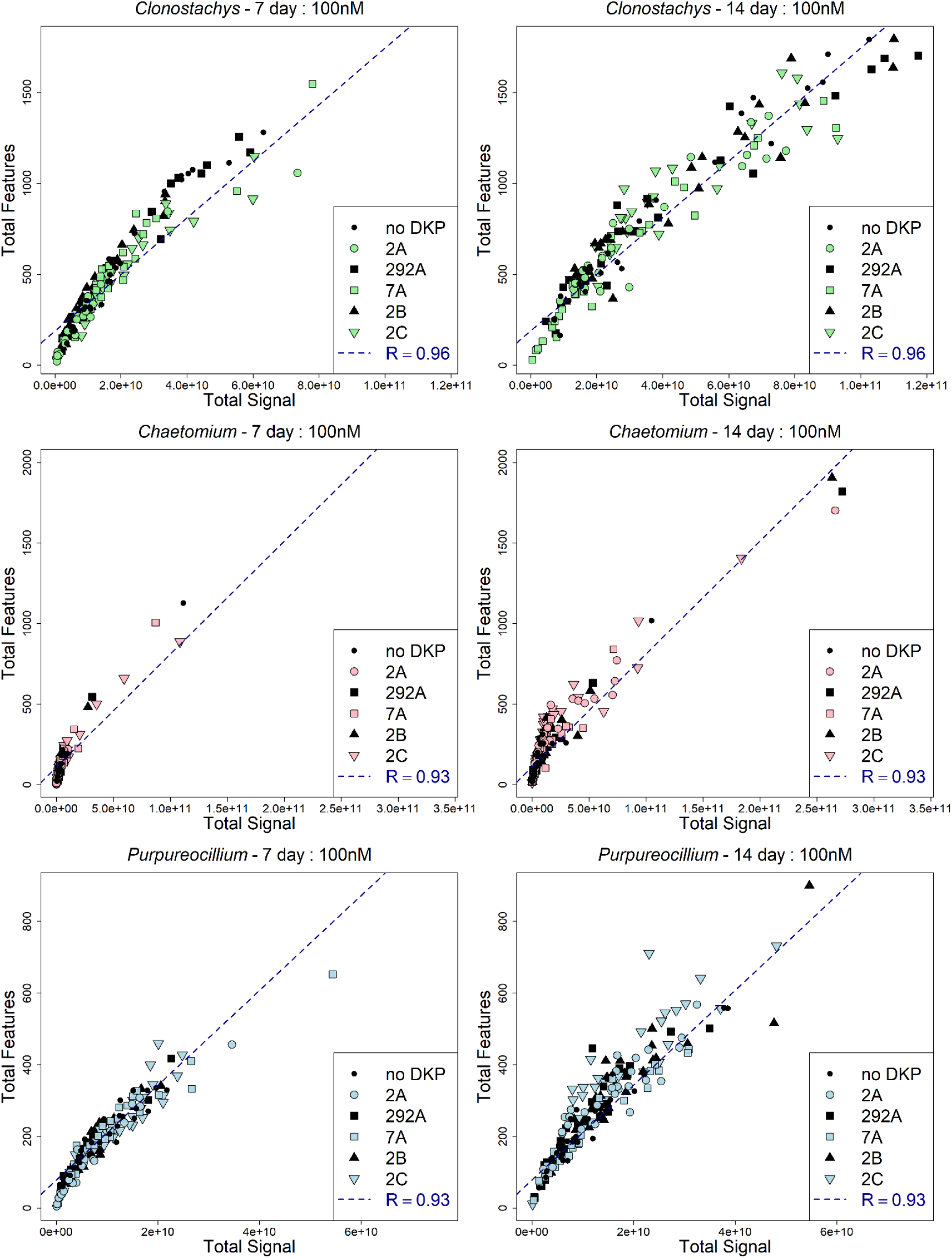
Total raw MS signal strongly correlates to chemical diversity - DKP 100 nM supplementation. For each timepoint and genera, a correlation between raw MS signal (after 5-fold blank and media feature removal)(See Data Processing) and total feature count was built. Raw MS signal was used as a proxy for biomass for this analysis.

## Tables

**Table S1.** Summary of average unique and total features per each region. This table shows results for the *Trichoderma* dataset.

| Region | Total Extracts | Total Features | Average Number of Features | Features Unique to Region | Average Unique Features per Extract |
| --- | --- | --- | --- | --- | --- |
| Alaska | 4 | 698 | 233 | 23 | 5.8 |
| Hawaii | 2 | 1392 | 724 | 51 | 25.5 |
| Northern Rockies | 7 | 3665 | 671 | 111 | 15.9 |
| Northwest | 41 | 10680 | 471 | 648 | 15.8 |
| West | 48 | 13222 | 519 | 1318 | 27.5 |
| Southwest | 35 | 14905 | 702 | 1633 | 46.7 |
| Upper Midwest | 70 | 22149 | 681 | 2187 | 31.2 |
| Northeast | 135 | 27925 | 666 | 2200 | 16.3 |
| Ohio Valley | 137 | 34132 | 741 | 3650 | 26.6 |
| Southeast | 194 | 36192 | 629 | 4831 | 24.9 |
| South | 160 | 39077 | 765 | 6317 | 39.5 |

**Table S2.** Summary of average unique and total features per each region. This table shows results for the multi-genera dataset.

| Region | Total Extracts | Total Features | Average Number of Features | Features Unique to Region | Average Unique Features per Extract |
| --- | --- | --- | --- | --- | --- |
| Alaska | 5 | 227 | 61 | 16 | 3.2 |
| Hawaii | 4 | 418 | 110 | 41 | 10.3 |
| Northern Rockies | 17 | 2251 | 168 | 281 | 16.5 |
| Northwest | 68 | 5261 | 119 | 878 | 12.9 |
| West | 87 | 7028 | 150 | 1536 | 17.7 |
| Southwest | 133 | 8378 | 120 | 1954 | 14.7 |
| Upper Midwest | 213 | 10067 | 106 | 1899 | 8.9 |
| Northeast | 194 | 10494 | 123 | 1953 | 10.1 |
| Ohio Valley | 212 | 12551 | 132 | 2579 | 12.2 |
| Southeast | 237 | 13136 | 124 | 2858 | 12.1 |
| South | 261 | 12831 | 114 | 3372 | 12.9 |

**Table S3.** Summary of average unique and total scaffolds per each region. This table shows results for the multi-genera dataset.

| Region | Total Extracts | Total Scaffolds | Average Number of Scaffolds | Scaffolds Unique to Region | Average Unique Scaffolds per Extract |
| --- | --- | --- | --- | --- | --- |
| Alaska | 5 | 136 | 40 | 0 | 0.0 |
| Hawaii | 4 | 198 | 57 | 1 | 0.3 |
| Northern Rockies | 17 | 707 | 75 | 7 | 0.4 |
| Northwest | 68 | 1442 | 57 | 36 | 0.5 |
| West | 87 | 1670 | 71 | 82 | 0.9 |
| Southwest | 133 | 1946 | 59 | 88 | 0.7 |
| Upper Midwest | 213 | 2233 | 53 | 61 | 0.3 |
| Northeast | 194 | 2315 | 61 | 70 | 0.4 |
| Ohio Valley | 212 | 2610 | 64 | 111 | 0.5 |
| Southeast | 237 | 2602 | 59 | 98 | 0.4 |
| South | 261 | 2642 | 56 | 153 | 0.6 |

**Table S4.** Summary of percent changes and standard errors, separated by fungal genus.

| Means | 7 Day |  |  | 14 Day |  |  |
| --- | --- | --- | --- | --- | --- | --- |
|  | *Clono-*<br>*stachys* | *Chaet-*<br>*omium* | *Purpur-*<br>*eocillium* | *Clono-*<br>*stachys* | *Chaet-*<br>*omium* | *Purpur-*<br>*eocillium* |
| Soil extract | 15% ± 7% | 40% ± 19% | 41% ± 10% | 4% ± 8% | -16% ± 16% | 19% ± 10% |
| Soil extract + sucrose | -28% ± 6% | -24% ± 12% | -22% ± 6% | 3% ± 10% | -22% ± 16% | -15% ± 9% |
| LPS - 0.06 ng/mL | -31% ± 5% | -40% ± 10% | -26% ± 6% | 5% ± 8% | -9% ± 17% | 12% ± 9% |
| LPS - 0.6 ng/mL | -33% ± 5% | -38% ± 10% | -34% ± 5% | 1% ± 8% | -11% ± 16% | -1% ± 9% |
| LPS - 1.8 ng/ml | -35% ± 5% | -41% ± 10% | -41% ± 6% | 16% ± 10% | -3% ± 20% | 4% ± 13% |
| DKP - 2A (10 µM) | -33% ± 10% | -38% ± 15% | -42% ± 10% | -30% ± 12% | -14% ± 22% | -40% ± 9% |
| DKP - 292A (10 µM) | -39% ± 9% | -26% ± 21% | -24% ± 8% | -9% ± 12% | 20% ± 30% | -17% ± 9% |
| DKP - 7A (10 µM) | -13% ± 13% | -7% ± 29% | -20% ± 10% | -28% ± 12% | -13% ± 32% | -40% ± 9% |
| DKP - 2B (10 µM) | -1% ± 14% | 11% ± 37% | 4% ± 14% | -20% ± 13% | -5% ± 22% | -29% ± 9% |
| DKP - 2C (10 µM) | 2% ± 13% | 62% ± 40% | 29% ± 12% | 1% ± 12% | 98% ± 38% | 49% ± 15% |
| DKP - Mixture (10 µM) | -38% ± 10% | -54% ± 13% | -41% ± 10% | -18% ± 10% | 77% ± 41% | 20% ± 12% |

**Table S5:** Mass Spectrometry Tune Data. Tune method data used by the Thermo Q-Exactive plus for all data acquisition.

| Parameter | Multi-gener<br>a | *Trichode*<br>*rma*<br>Extracts | Soils/LP<br>S<br>Addition | Diketopi<br>perazine<br>Addition |
| --- | --- | --- | --- | --- |
| **Instrument** | Q-Exactive<br>Plus | Q-Exacti<br>ve Plus | Q-Exacti<br>ve Plus | Q-Exactiv<br>e Plus |
| **Injection Volume (µL)** | 5 | 5 | 5 | 5 |
| **Tune Method** |  |  |  |  |
| Spray Voltage (+, V) | 3800.00 | 3800.00 | 3800.00 | 3800.00 |
| Capillary Temperature (°C) | 320.00 | 320.00 | 320.00 | 320.00 |
| Sheath Gas (arbitrary units) | 35.00 | 35.00 | 35.00 | 35.00 |
| Aux Gas (arbitrary units) | 10.00 | 10.00 | 10.00 | 10.00 |
| Spare Gas (arbitrary units) | 0.00 | 0.00 | 0.00 | 0.00 |
| Max Spray Current (µA) | 100.00 | 100.00 | 100.00 | 100.00 |
| Probe Heater Temp (°C) | 350.00 | 350.00 | 350.00 | 350.00 |
| S-Lens RF Level (Thermo Arbitrary units) | 50.00 | 50.00 | 50.00 | 50.00 |
| Ion Source | HESI | HESI | HESI | HESI |

**Table S6:** MS/MS Data Acquisition Parameters.

| Parameter | Multi-gener<br>a collection | *Trichode*<br>*rma*<br>Extracts | Soils/LP<br>S<br>Addition | Diketopi<br>perazine<br>Addition |
| --- | --- | --- | --- | --- |
| **General Experiment** |  |  |  |  |
| Type | Full MS/<br>DDMS2<br>(TopN) | Full MS/<br>DDMS2<br>(TopN) | Full MS/<br>DDMS2<br>(TopN) | Full MS/<br>DDMS2<br>(TopN) |
| Runtime (min) | 0-12.5 | 0-12.5 | 0-12.5 | 0-12.5 |
| Polarity | positive | positive | positive | positive |
| In-source CID (eV) | 0.0 | 0.0 | 0.0 | 0.0 |
| Default Charge State | 1 | 1 | 1 | 1 |
| Exclusion | on | on | on | on |
| Exclusion Description | Top 500 in<br>blank | Top 3000<br>in blank | Top 3000<br>in blank | Top 3000<br>in blank |
| **Full MS** |  |  |  |  |
| Microscan | 1 | 1 | 1 | 1 |
| Resolution | 70,000 | 70,000 | 70,000 | 70,000 |
| AGC target | 1e6 | 1e6 | 1e6 | 1e6 |
| Maximum IT (ms) | 246 | 246 | 246 | 246 |
| Number of Scan ranges | 1 | 1 | 1 | 1 |
| Scan Range | *m/z*<br>100-1500 | *m/z*<br>120-1200 | *m/z*<br>120-1200 | *m/z*<br>70-1050 |
| Spectrum data Type | profile | profile | profile | profile |
| **dd-MS2** |  |  |  |  |
| microscans | 1 | 1 | 1 | 1 |
| Resolution | 17,500 | 17,500 | 17,500 | 17,500 |
| AGC target | 2e5 | 2e5 | 2e5 | 2e5 |
| Maximum IT (ms) | 54 | 54 | 54 | 54 |
| Loop count | 5 | 5 | 5 | 5 |
| MSX count | 1 | 1 | 1 | 1 |
| TopN | 5 | 5 | 5 | 5 |
| Isolation Window ($m/z$) | 1.0 | 1.0 | 1.0 | 1.0 |
| Isolation offset ($m/z$) | 0 | 0 | 0 | 0 |
| Scan range ($m/z$) | 200-2000 | 200-2000 | 200-2000 | 200-2000 |
| Fixed first mass | — | — | — | — |
| NCE | 20,40,60 | 20,40,60 | 20,40,60 | 20,40,60 |
| **dd settings** |  |  |  |  |
| Minimum AGC target | 8.00e3 | 8.00e3 | 8.00e3 | 8.00e3 |
| Intensity threshold (AU) | 1.5e5 | 1.5e5 | 1.5e5 | 1.5e5 |
| Apex trigger | — | — | — | — |
| Charge exclusion | — | — | — | — |
| Multiple charge states | all | all | all | all |
| Peptide match | preferred | preferred | preferred | preferred |
| Exclude isotopes | on | on | on | on |
| Dynamic exclusion (s) | 10.0 | 10.0 | 10.0 | 10.0 |
| If idle | Do not pick others | Do not pick others | Do not pick others | Do not pick others |
| **Misc** |  |  |  |  |
| Divert valve goes to MS (min) | 0.20-12.5 | 0.20-12.5 | 0.20-12.5 | 0.20-12.5 |
| Inclusion list | none | none | none | none |

**Table S7:**
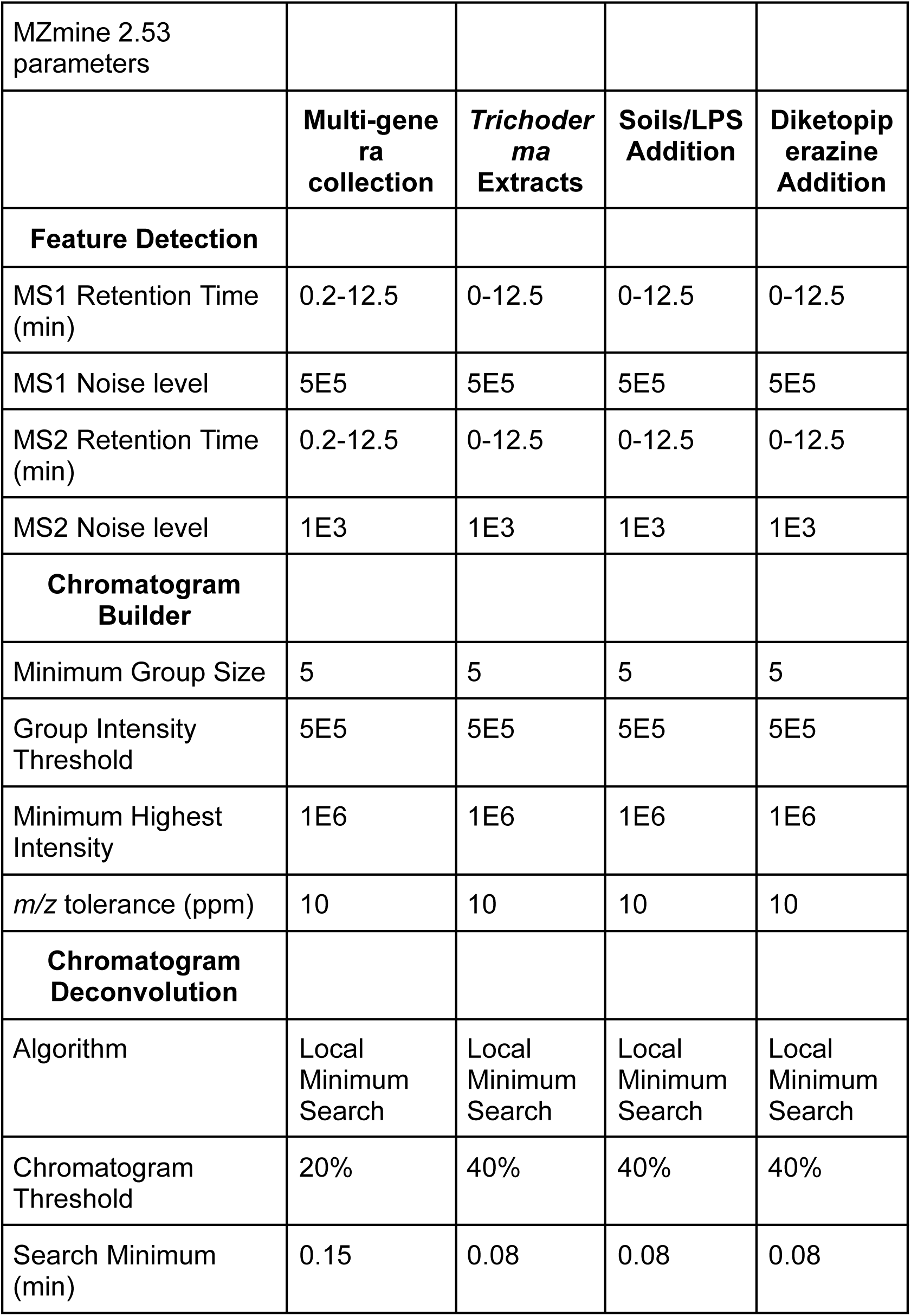
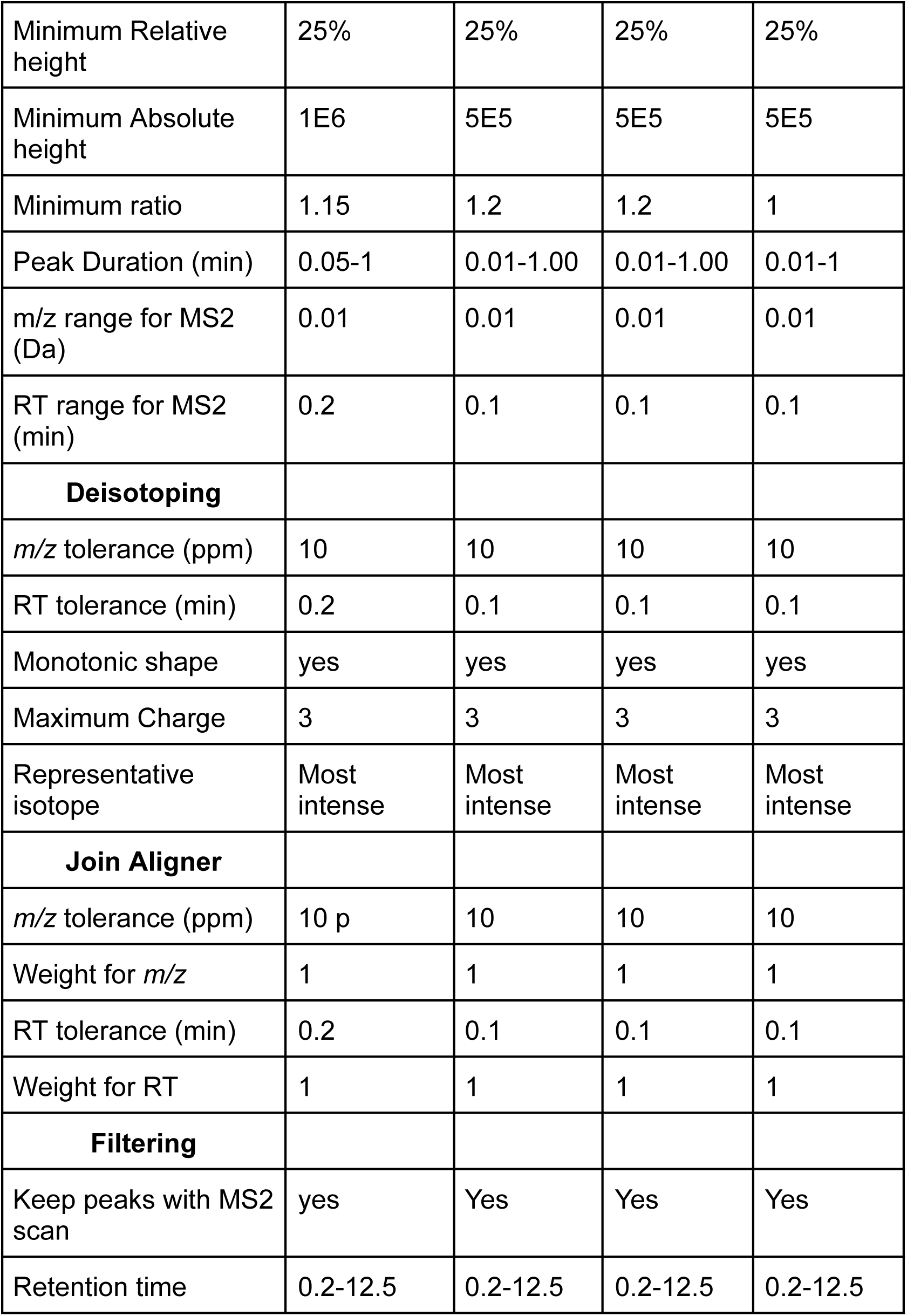
MZmine 2.53 parameters. MZmine parameters used for all runs. Parameters for the multi-genera collection taken from a previous analysis ^17^.

|  |  |  |  |  |
| --- | --- | --- | --- | --- |
| MZmine 2.53 parameters |  |  |  |  |
|  | **Multi-genera collection** | ***Trichoderma* Extracts** | **Soils/LPS Addition** | **Diketopiperazine Addition** |
| **Feature Detection** |  |  |  |  |
| MS1 Retention Time (min) | 0.2-12.5 | 0-12.5 | 0-12.5 | 0-12.5 |
| MS1 Noise level | 5E5 | 5E5 | 5E5 | 5E5 |
| MS2 Retention Time (min) | 0.2-12.5 | 0-12.5 | 0-12.5 | 0-12.5 |
| MS2 Noise level | 1E3 | 1E3 | 1E3 | 1E3 |
| **Chromatogram Builder** |  |  |  |  |
| Minimum Group Size | 5 | 5 | 5 | 5 |
| Group Intensity Threshold | 5E5 | 5E5 | 5E5 | 5E5 |
| Minimum Highest Intensity | 1E6 | 1E6 | 1E6 | 1E6 |
| *m/z* tolerance (ppm) | 10 | 10 | 10 | 10 |
| **Chromatogram Deconvolution** |  |  |  |  |
| Algorithm | Local Minimum Search | Local Minimum Search | Local Minimum Search | Local Minimum Search |
| Chromatogram Threshold | 20% | 40% | 40% | 40% |
| Search Minimum (min) | 0.15 | 0.08 | 0.08 | 0.08 |
| Minimum Relative height | 25% | 25% | 25% | 25% |
| Minimum Absolute height | 1E6 | 5E5 | 5E5 | 5E5 |
| Minimum ratio | 1.15 | 1.2 | 1.2 | 1 |
| Peak Duration (min) | 0.05-1 | 0.01-1.00 | 0.01-1.00 | 0.01-1 |
| m/z range for MS2 (Da) | 0.01 | 0.01 | 0.01 | 0.01 |
| RT range for MS2 (min) | 0.2 | 0.1 | 0.1 | 0.1 |
| **Deisotoping** |  |  |  |  |
| *m/z* tolerance (ppm) | 10 | 10 | 10 | 10 |
| RT tolerance (min) | 0.2 | 0.1 | 0.1 | 0.1 |
| Monotonic shape | yes | yes | yes | yes |
| Maximum Charge | 3 | 3 | 3 | 3 |
| Representative isotope | Most intense | Most intense | Most intense | Most intense |
| **Join Aligner** |  |  |  |  |
| *m/z* tolerance (ppm) | 10 p | 10 | 10 | 10 |
| Weight for *m/z* | 1 | 1 | 1 | 1 |
| RT tolerance (min) | 0.2 | 0.1 | 0.1 | 0.1 |
| Weight for RT | 1 | 1 | 1 | 1 |
| **Filtering** |  |  |  |  |
| Keep peaks with MS2 scan | yes | Yes | Yes | Yes |
| Retention time | 0.2-12.5 | 0.2-12.5 | 0.2-12.5 | 0.2-12.5 |

**Table S8:** GNPS feature-based molecular networking parameters.

|  |  |
| --- | --- |
| **Basic Options** |  |
| Table Source | MZmine |
| Ion Mass Tolerance | 0.02 Da |
| Fragment Ion Mass Tolerance | 0.02 Da |
| **Advanced Network Options** |  |
| Min Pairs Cos | 0.7 |
| Network TopK | 10 |
| Minimum Matched Fragment Ions | 4 |
| Maximum Connected Component Size | 100 |
| Maximum shift between precursors | 500 Da |
| **Advanced Library Search Options** |  |
| Spectral Library | Shown in Table S9 |
| Library Search Min Max Peaks | 6 |
| Search Analogs | Don't Search |
| Top Results to Report | 1 |
| Score Threshold | 0.7 |
| Maximum Analog Search Mass Difference | 100 Da |
| **Advanced Filtering Options** |  |
| Minimum Peak Intensity | 0 |
| Filter Precursor Window | Filter |
| Filter Peaks in 50 Da Window | Filter |
| Filter Library | Filter library |
| **Advanced Quantification Options** |  |
| Normalization per File | Row Sum<br>Normalization |
| Aggregation Method | Mean |
| **Advanced Multivariable**<br>**Statistics Options** |  |
| PCoA Distance Metric | Cosine |
| **Advanced Univariate Statistics**<br>**Options** |  |
| Metadata Column to Compare | None |
| Metadata Field to Compare | None |
| Metadata Column to Facet | None |
| Run Stats and Plots | No |
| Metadata Field to Compare | None |
| **Advanced External Tools** |  |
| Run Dereplicator | Run |

**Table S9:** GNPS Spectral Libraries. . Spectral libraries used for feature-based molecular networking spectral matching.

|  |  |  |
| --- | --- | --- |
| BERKLEY-LAB | GNPS-NIH-SMALLMOLECULEPHARMACOLOGICALYACTIVE | MIADB |
| BMDMS-NP | GNPS-NIST14-MATCHES | MMVPOSITIVE |
| DEREPLICATOR_IDENTIFIED_LIBRARY | GNPS-PRESTWICKPHYTOCHEM | MONA |
| ECG-ACYL-AMIDES-C4-C24-LIBRARY | GNPS-SAM-SIK-KANG-LEGANCY-LIBRARY | PNNL-LIPIDS |
| ECG-ACYL-ESTERS-C4-C24-LIBRARY | GNPS-SELLECKCHEM-FDA-PART1 | RESPECT |
| GNPS-COLLECTIONS-MISC | GNPS-SELLECKCHEM-FDA-PART2 | SUMNER |
| GNPS-COLLECTIONS-PESTICIDES-POSITIVE | LDB_POSITIVE | UM-NPDC |
| GNPS-D2-AMINO-LIPID-LIBRARY | MASSBANK | XANTHONES-DB |
| GNPS-NIH-NATURALPRODUCTSLIBRARY_ROUND2_POSITIVE | MASSBANKEU |  |

