## Supplemental Data 1 for "Systematic assessment of the impact of targeted selection methods and environment-mimicking culture conditions on fungal natural product libraries"

<u>**Supplemental Data 1: Synthesis of diketopiperazines**</u>

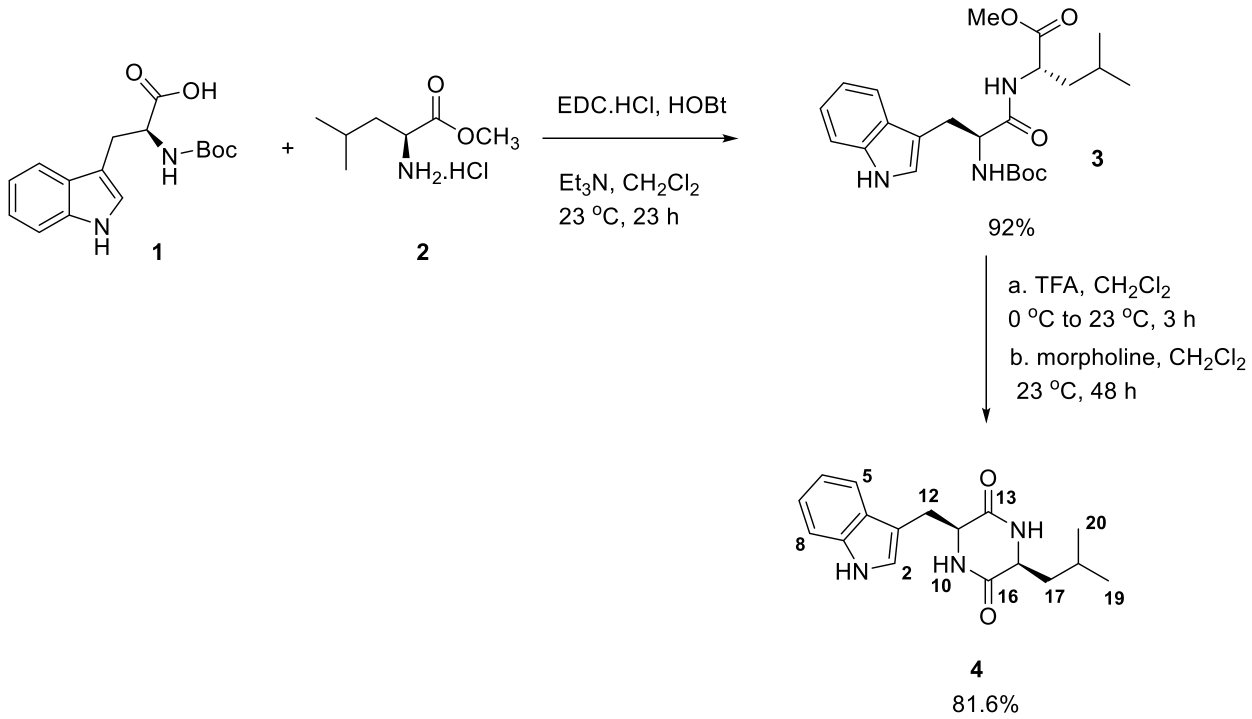


(tert-butoxycarbonyl)-L-tryptophan **1** (1 g, 3.3 mmol, 1 equiv), L-leucine methyl ester hydrochloride **2** (0.8 g, 4.9 mmol, 1.5 equiv), and 1-hydroxybenzotriazole hydrate (0.6 g, 4.9 mmol, 1.5 equiv) were suspended in dichloromethane (50 mL). Triethylamine (2 mL, 14.7 mmol, 4.5 equiv) was added to the mixture and the suspension was stirred until a golden homogenous solution was formed (5-10 min). To this solution EDC.HCl (0.9 g, 4.92 mmol, 1.5 equiv) was added and the resulting mixture was stirred for 8 h before another portion of EDC.HCl (0.3 g, 1.96 mmol, 0.6 equiv) and triethylamine (0.8 mL, 6.56 mmol, 2 equiv) were added. The reaction mixture was stirred for an additional 15 h. The mixture was first washed with an aqueous solution of HCl (1N, 20 mL) and saturated aqueous $NaHCO_3$ solution (20 mL). The organic layer was dried over anhydrous $Na_2SO_4$, filtered and was concentrated in vacuo to afford a yellow foam. The crude was purified using a silica column (30% ethyl acetate in hexanes) to yield the product as a white foam **3** (1.3 g, 92%).

The resulting dipeptide (1.3 g, 3.28 mmol, 1 equiv) was dissolved in dichloromethane (10 mL) and trifluoracetic acid (3.0 mL) was added at 0 °C. The volatiles were removed after 3 h and the residue was redissolved in dichloromethane (50 mL). To the resulting mixture morpholine (8 mL) was added and the solution was vigorously stirred for 48 h as the product was precipitated as a white solid. The product was filtered under reduced pressure and washed with DCM (3 × 30 mL) and deionized water (3 × 30 mL) to afford **4** as a white solid (0.80 g, 81.6%). The product was further purified using preparative HPLC, C18 column using 80% ACN/$H_2O$ with 0.1% FA.

$^1$H NMR (600 MHz, DMSO-$d_6$): δ10.89 (s, 1H, $N_1$**H**), 7.02 (s, 1H, $C_2$**H**), 7.55 (d, *J* = 7.8 Hz, 1H, $C_5$**H**), 6.92 (t, *J* = 7.2 Hz, *J* = 1.2 Hz, 1H, $C_6$**H**), 7.02 (t, *J* = 7.2 Hz, 1H, $C_7$**H**), 7.30 (d, *J* = 7.8 Hz, 1H, $C_8$**H**), 8.02 (brd, *J* = 2.4 Hz, 1H, $N_{10}$**H**), 4.09 (q, *J* = 3.6 Hz ,1H, $C_{11}$**H**), 3.25 (dd, *J* = 14.4 Hz, 4.2 Hz, 1H, $C_{12}$**H**), 2.89 (dd, *J* = 14.4 Hz, 4.8 Hz, 1H, $C_{12}$**H**), 7.93 (brd, *J* = 3.0,1H, $N_{14}$**H**), 3.40 (m, 1H, $C_{15}$**H**), 2.55 (m, 1H, $C_{17}$**H**), 0.60 (m, 1H, $C_{17}$**H**), 1.21 (sept, *J* = 6.6 Hz, 1H, $C_{18}$**H**), 3.0 (d, *J* = 6.6 Hz, 3H, $C_{19}$**H),** 2.97 (d, *J* = 6.6 Hz, 3H, $C_{20}$**H)**

$^{13}$C NMR (100 MHz, DMSO-$d_6$): δ124.7(C**2**), 108.5 (C**3**), 127.8 (C**4**), 118.4 (C**5**), 119.0 (C**6**), 120.8 (C**7**), 111.2 (C**8**), 136.0 (C**9**), 52.4 (C**11**), 29.1 (C**12**), 167.5 (C**13**), 55.6 (C**15**), 167.2 (C**16**), 43.7 (C**17**), 22.9 (C**18**), 22.7 (C**19**), 21.3 (C**20**)

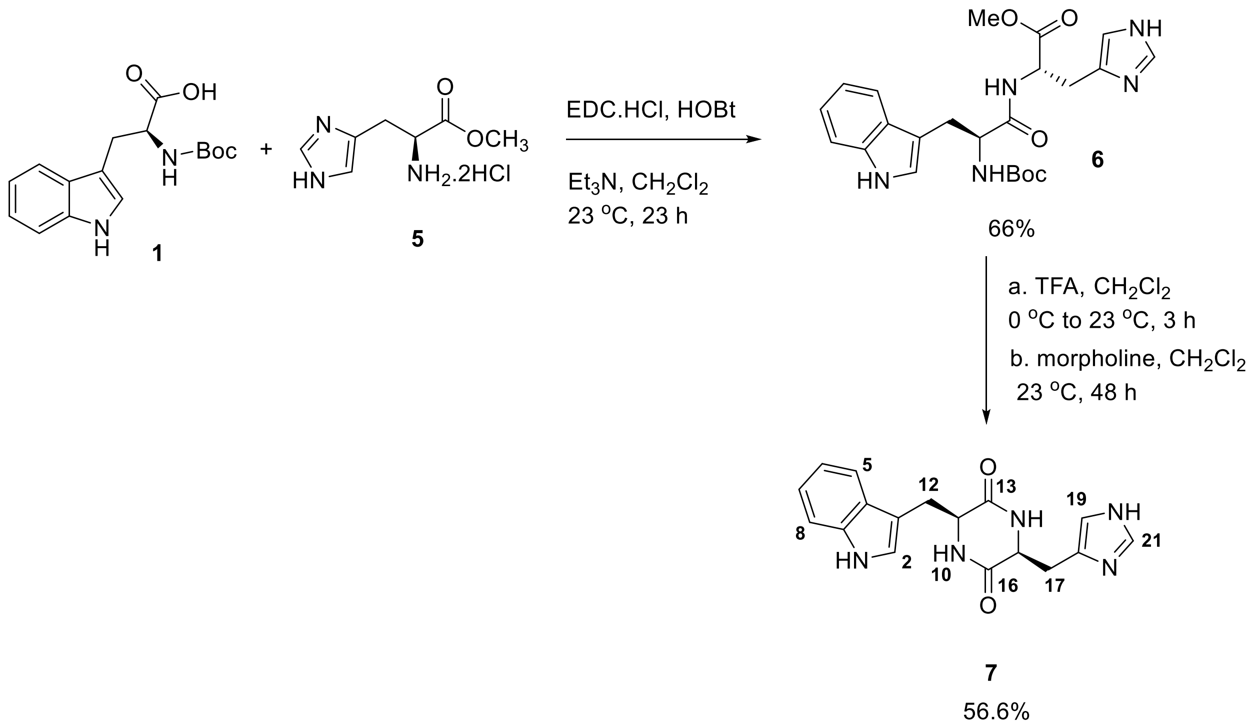

(tert-butoxycarbonyl)-L-tryptophan **1** (1 g, 3.3 mmol, 1 equiv), L-histidine methyl ester dihydrochloride **5** (1.2 g, 4.9 mmol, 1.5 equiv), and 1-hydroxybenzotriazole hydrate (0.6 g, 4.9 mmol, 1.5 equiv) were suspended in dichloromethane (50 mL). Triethylamine (2 mL, 14.7 mmol, 4.5 equiv) was added to the mixture and the suspension was stirred until a golden homogenous solution was formed (5-10 min). To this solution EDC.HCl (0.9 g, 4.92 mmol, 1.5 equiv) was added and the resulting mixture was stirred for 8 h before another portion of EDC.HCl (0.3 g, 1.96 mmol, 0.6 equiv) and triethylamine (0.8 mL, 6.56 mmol, 2 equiv) were added. The reaction mixture was stirred for an additional 15 h. The mixture was first washed with an aqueous solution of HCl (1N, 20 mL) and saturated aqueous $NaHCO_3$ solution (20 mL). The organic layer was dried over anhydrous $Na_2SO_4$, filtered and was concentrated in vacuo to afford a yellow foam. The crude was purified using a silica column (42% ethyl acetate in hexanes) to yield the product as a white foam **6** (1.0 g, 66%).
The resulting dipeptide (1.0 g, 3.28 mmol, 1 equiv) was dissolved in dichloromethane (10 mL) and trifluoracetic acid (3.0 mL) was added at 0 °C. The volatiles were removed after 3 h and the residue was redissolved in dichloromethane (50 mL). To the resulting mixture morpholine (8 mL) was added and the solution was vigorously stirred for 48 h as the product was precipitated as a white solid. The product was filtered under reduced pressure and washed with DCM (3 × 30 mL) and deionized water (3 × 30 mL) to afford **7** as a white solid (0.60 g, 56.6%). The product was further purified using preparative HPLC, C18 column using 70% ACN/$H_2O$ with 0.1% FA.

$^1$H NMR (600 MHz, DMSO-$d_6$): δ10.90 (s, 1H, $N_1$**H**), 7.05 (s, 1H, $C_2$**H**), 7.54 (d, $J$ = 7.8 Hz, 1H, $C_5$**H**), 6.95 (t, $J$ = 7.8 Hz, 1H, $C_6$**H**), 7.04 (t, $J$ = 7.8 Hz, 1H, $C_7$**H**), 7.32 (d, $J$ = 7.8 Hz, 1H, $C_8$**H**), 7.57 (brd, $J$ = 2.4 Hz, 1H, $N_{10}$**H**), 4.07 (m,1H, $C_{11}$**H**), 3.03 (dd, $J$ = 14.4 Hz, 4.8 Hz, 1H, $C_{12}$**H**), 2.96 (dd, $J$ = 14.4 Hz, 4.2 Hz, 1H, $C_{12}$**H**), 8.01 (brd, $J$ = 2.4,1H, $N_{14}$**H**), 3.74 (m, 1H, $C_{15}$**H**), 2.39 (dd, $J$ = 14.4 Hz, 4.2 Hz, 1H, $C_{17}$**H**), 1.30 (dd, $J$ = 14.4 Hz, 4.8 Hz, 1H, $C_{17}$**H**), 7.68 (brs, 1H, $C_{19}$**H),** 8.12 (brs, 1H, $C_{21}$**H)**

$^{13}$C NMR (100 MHz, DMSO-$d_6$): δ124.7(C**2**), 108.6 (C**3**), 127.7 (C**4**), 120.8 (C**5**), 118.4 (C**6**), 119.0 (C**7**), 111.3 (C**8**), 134.8 (C**9**), 54.2 (C**11**), 29.0 (C**12**), 166.4 (C**13**), 55.3 (C**15**), 166.9 (C**16**), 31.0 (C**17**), 131.8 (C**18**), 117.1 (C**19**), 135.9 (C**21**)

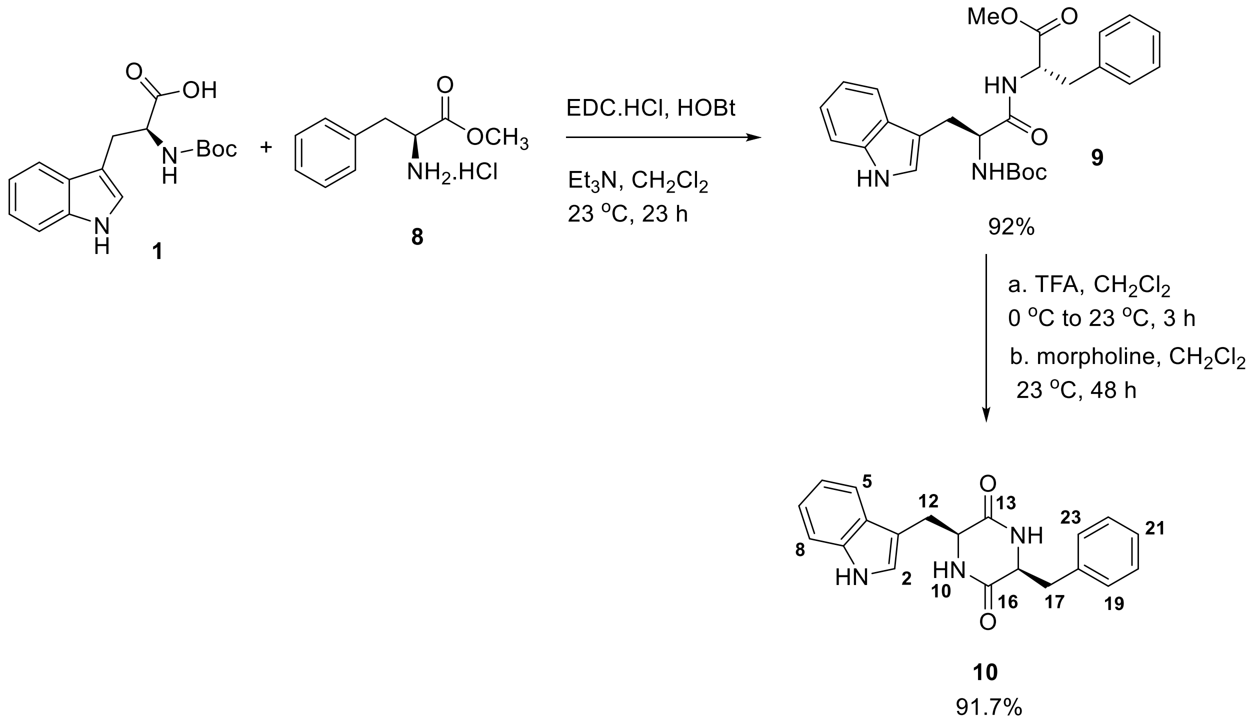


(tert-butoxycarbonyl)-L-tryptophan **1** (1 g, 3.3 mmol, 1 equiv), L-phenylalanine methyl ester hydrochloride **8** (0.9 g, 4.9 mmol, 1.5 equiv), and 1-hydroxybenzotriazole hydrate (0.6 g, 4.9 mmol, 1.5 equiv) were suspended in dichloromethane (50 mL). Triethylamine (2 mL, 14.7 mmol, 4.5 equiv) was added to the mixture and the suspension was stirred until a golden homogenous solution was formed (5-10 min). To this solution EDC.HCl (0.9 g, 4.92 mmol, 1.5 equiv) was added and the resulting mixture was stirred for 8 h before another portion of EDC.HCl (0.3 g, 1.96 mmol, 0.6 equiv) and triethylamine (0.8 mL, 6.56 mmol, 2 equiv) were added. The reaction mixture was stirred for an additional 15 h. The mixture was first washed with an aqueous solution of HCl (1N, 20 mL) and saturated aqueous $NaHCO_3$ solution (20 mL). The organic layer was dried over anhydrous $Na_2SO_4$, filtered and was concentrated in vacuo to afford a yellow foam. The crude was purified using a silica column (25% ethyl acetate in hexanes) to yield the product as a white foam **9** (1.4 g, 92%).

The resulting dipeptide (1.4 g, 3.28 mmol, 1 equiv) was dissolved in dichloromethane (10 mL) and trifluoracetic acid (3.0 mL) was added at 0 °C. The volatiles were removed after 3 h and the residue was redissolved in dichloromethane (50 mL). To the resulting mixture morpholine (8 mL) was added and the solution was vigorously stirred for 48 h as the product was precipitated as a white solid. The product was filtered under reduced pressure and washed with DCM (3 × 30 mL) and deionized water (3 × 30 mL) to afford **10** as a white solid (1.0 g, 91.7%). The product was further purified using preparative HPLC, C18 column using 80% ACN/$H_2O$ with 0.1% FA.

$^1H$ NMR (400 MHz, DMSO-$d_6$): δ10.88 (s, 1H, $N_1$**H**), 6.96 (s, 1H, $C_2$**H**), 7.32 (dt, *J* = 8.0 Hz, 1.2 Hz, 1H, $C_5$**H**), 6.98 (td, *J* = 7.2 Hz, *J* = 1.2 Hz, 1H, $C_6$**H**), 7.06 (td, *J* = 7.2 Hz, *J* = 1.2 Hz, 1H, $C_7$**H**), 7.48 (d, *J* = 8.0 Hz, 1H, $C_8$**H**), 7.69 (brd, *J* = 2.8 Hz, 1H, $N_{10}$**H**), 3.85 (m, 1H, $C_{11}$**H**), 2.80 (dd, *J* = 14.4 Hz, 4.8 Hz, 1H, $C_{12}$**H**), 2.46 (dd, *J* = 13.6 Hz, 4.8 Hz, 1H, $C_{12}$**H**), 7.88 (brd, *J* = 2.8,1H, $N_{14}$**H**), 3.97 (m, 1H, $C_{15}$**H**), 2.55 (dd, *J* = 13.3 Hz, 4.8 Hz, 1H, $C_{17}$**H**), 2.45 (dd, *J* = 13.3 Hz, 4.8 Hz, 1H, $C_{17}$**H**), 6.71 (dd, *J* = 7.6 Hz, 2.0 Hz, 2H, $C_{19}$**H**, $C_{23}$**H**), 7.14 – 7.20 (m, 3H, $C_{20}$**H**, $C_{21}$**H**, $C_{22}$**H**)

$^{13}C$ NMR (100 MHz, DMSO-$d_6$): δ126.3 (C**2**), 108.8 (C**3**), 124.3 (C**4**), 120.8 (C**5**), 118.7 (C**6**), 118.3 (C**7**), 111.3 (C**8**), 136.0 (C**9**), 55.6 (C**11**), 29.6 (C**12**), 166.7 (C**13**), 55.2 (C**15**), 166.1 (C**16**), 40.1 (C**17**), 136.5 (C**18**), 128.0 (C**19**, C**23**), 129.6 (C**20**, C**22**), 127.5 (C**21**)

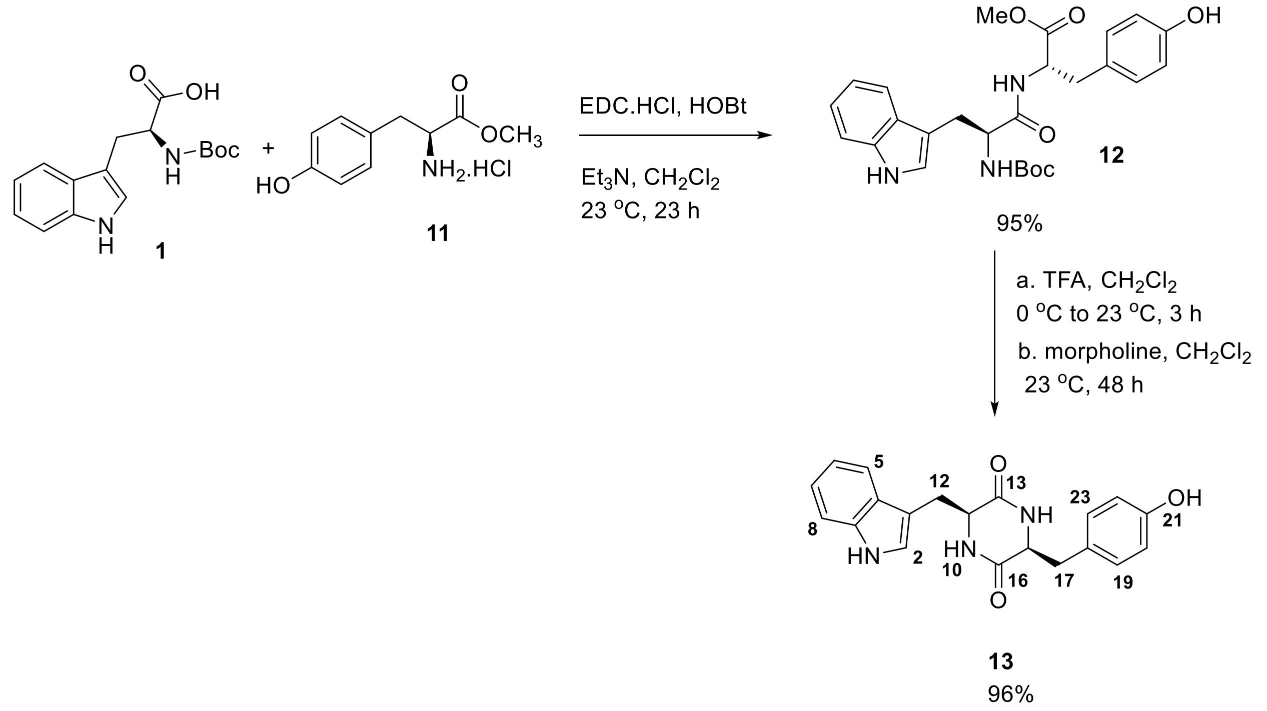


(tert-butoxycarbonyl)-L-tryptophan **1** (1 g, 3.3 mmol*,* 1 equiv), L-tyrosine methyl ester hydrochloride **11** (1.1 g, 4.9 mmol, 1.5 equiv), and 1-hydroxybenzotriazole hydrate (0.6 g, 4.9 mmol, 1.5 equiv) were suspended in dichloromethane (50 mL). Triethylamine (2 mL, 14.7

mmol, 4.5 equiv) was added to the mixture and the suspension was stirred until a golden homogenous solution was formed (5-10 min). To this solution EDC.HCl (0.9 g, 4.92 mmol, 1.5 equiv) was added and the resulting mixture was stirred for 8 h before another portion of EDC.HCl (0.3 g, 1.96 mmol, 0.6 equiv) and triethylamine (0.8 mL, 6.56 mmol, 2 equiv) were added. The reaction mixture was stirred for an additional 15 h. The mixture was first washed with an aqueous solution of HCl (1N, 20 mL) and saturated aqueous $NaHCO_3$ solution (20 mL). The organic layer was dried over anhydrous $Na_2SO_4$, filtered and was concentrated in vacuo to afford a yellow foam. The crude was purified using a silica column (40% ethyl acetate in hexanes) to yield the product as a white foam **12** (1.5 g, 95%).

The resulting dipeptide (1.5 g, 3.28 mmol, 1 equiv) was dissolved in dichloromethane (10 mL) and trifluoracetic acid (3.0 mL) was added at 0 °C. The volatiles were removed after 3 h and the residue was redissolved in dichloromethane (50 mL). To the resulting mixture morpholine (8 mL) was added and the solution was vigorously stirred for 48 h as the product was precipitated as a white solid. The product was filtered under reduced pressure and washed with DCM (3 × 30 mL) and deionized water (3 × 30 mL) to afford **13** as a white solid (1.1 g, 96%). The product was further purified using preparative HPLC, C18 column using 70% ACN/$H_2O$ with 0.1% FA.

$^1$H NMR (400 MHz, DMSO-$d_6$): δ10.88 (s, 1H, $N_1$**H**), 6.97 (s, 1H, $C_2$**H**), 7.31 (dt, $J$ = 8.4 Hz, 1.2 Hz, 1H, $C_5$**H**), 6.98 (td, $J$ = 6.4 Hz, $J$ = 1.2 Hz, 1H, $C_6$**H**), 7.06 (td, $J$ = 6.6 Hz, $J$ = 1.2 Hz, 1H, $C_7$**H**), 7.47 (d, $J$ = 7.8 Hz, 1H, $C_8$**H**), 7.61 (brd, $J$ = 3.0 Hz, 1H, $N_{10}$**H**), 3.77 (m, 1H, $C_{11}$**H**), 2.80 (dd, $J$ = 14.4 Hz, 4.2 Hz, 1H, $C_{12}$**H**), 2.46 (dd, $J$ = 14.4 Hz, 6.0 Hz, 1H, $C_{12}$**H**), 7.79 (brd, $J$ = 3.0,1H, $N_{14}$**H**), 3.93 (m, 1H, $C_{15}$**H**), 2.43 (dd, $J$ = 14.4 Hz, 4.8 Hz, 1H, $C_{17}$**H**), 2.41 (dd, $J$ = 14.4 Hz, 4.8 Hz, 1H, $C_{17}$**H**), 6.58 (dd, $J$ = 8.4 Hz, 2H, $C_{19}$**H**, $C_{23}$**H**), 6.53 (dd, $J$ = 8.4 Hz, 2H, $C_{20}$**H**, $C_{22}$**H**), 9.14 (s, 1H, O**H**)

$^{13}$C NMR (100 MHz, DMSO-$d_6$): δ124.3 (C**2**), 108.9 (C**3**), 126.4 (C**4**), 118.4 (C**5**), 118.7 (C**6**), 120.8 (C**7**), 111.3 (C**8**), 136.0 (C**9**), 55.2 (C**11**), 29.9 (C**12**), 166.7 (C**13**), 55.8 (C**15**), 166.2 (C**16**), 38.9 (C**17**), 127.4 (C**18**), 130.7 (C**19**, C**23**), 114.8 (C**20**, C**22**), 155.9 (C**21**)

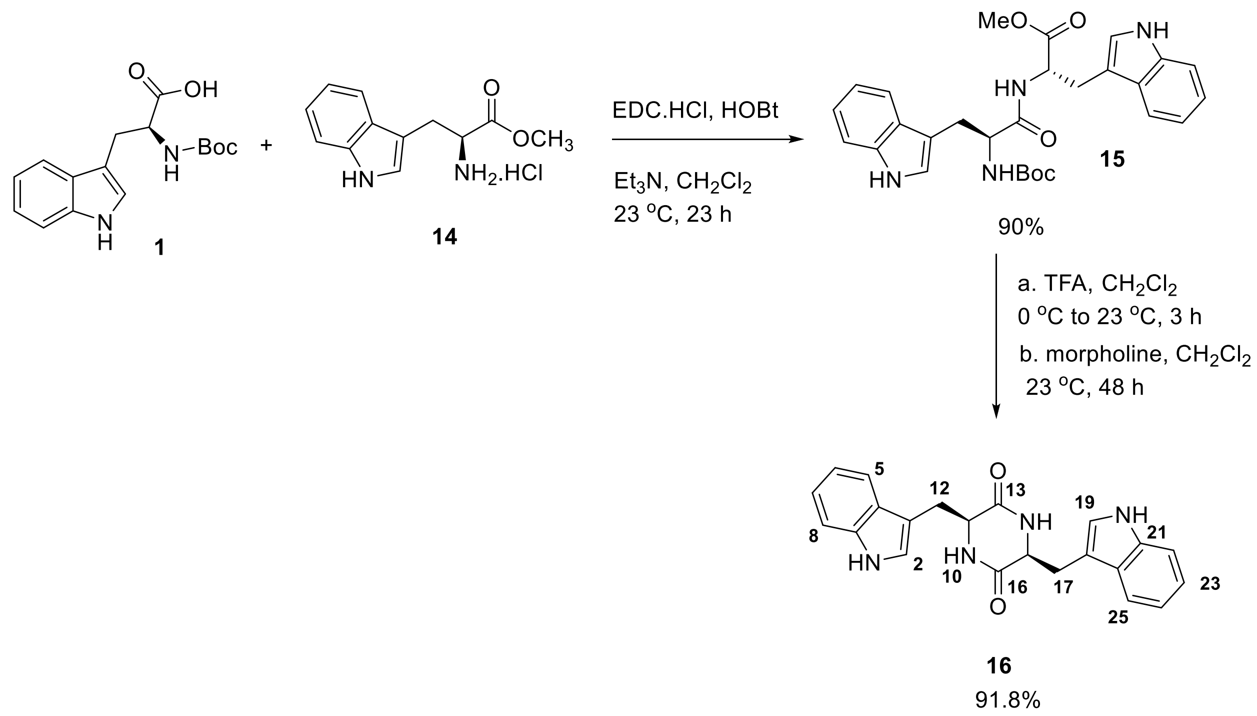


(tert-butoxycarbonyl)-L-tryptophan **1** (1 g, 3.3 mmol*,* 1 equiv), L-tyrosine methyl ester hydrochloride **14** (0.8 g, 4.9 mmol, 1.5 equiv), and 1-hydroxybenzotriazole hydrate (0.6 g, 4.9 mmol, 1.5 equiv) were suspended in dichloromethane (50 mL). Triethylamine (2 mL, 14.7 mmol, 4.5 equiv) was added to the mixture and the suspension was stirred until a golden homogenous solution was formed (5-10 min). To this solution EDC.HCl (0.9 g, 4.92 mmol, 1.5 equiv) was added and the resulting mixture was stirred for 8 h before another portion of EDC.HCl (0.3 g, 1.96 mmol, 0.6 equiv) and triethylamine (0.8 mL, 6.56 mmol, 2 equiv) were added. The reaction mixture was stirred for an additional 15 h. The mixture was first washed with an aqueous solution of HCl (1N, 20 mL) and saturated aqueous $NaHCO_3$ solution (20 mL). The organic layer was dried over anhydrous $Na_2SO_4$, filtered and was concentrated in vacuo to afford a yellow foam. The crude was purified using a silica column (25% ethyl acetate in hexanes) to yield the product as a white foam **15** (1.5 g, 90%).

The resulting dipeptide (1.5 g, 3.28 mmol, 1 equiv) was dissolved in dichloromethane (10 mL) and trifluoracetic acid (3.0 mL) was added at 0 °C. The volatiles were removed after 3 h and the residue was redissolved in dichloromethane (50 mL). To the resulting mixture morpholine (8 mL) was added and the solution was vigorously stirred for 48 h as the product was precipitated as a white solid. The product was filtered under reduced pressure and washed with DCM (3 × 30 mL) and deionized water (3 × 30 mL) to afford **16** as a white solid (1.1 g, 91.8%). The product was further purified using preparative HPLC, C18 column using 80% ACN/$H_2O$ with 0.1% FA.

$^{1}$H NMR (600 MHz, DMSO-*d*$_6$): δ10.81 (brd, *J* = 2.4 Hz, 2H, N$_1$**H**, N$_{20}$**H**), 6.60 (brd, *J* = 2.4 Hz, 2H, C$_2$**H**, C$_{19}$**H**), 7.34 (d, *J* = 7.8 Hz, 2H, C$_5$**H**, C$_{25}$**H**), 6.95 (td, *J* = 8.4 Hz, *J* = 1.2 Hz, 2H, C$_6$**H**, C$_{24}$**H**), 7.04 (td, *J* = 7.8 Hz, *J* = 1.2 Hz, 2H, C$_7$**H**, C$_{23}$**H**), 7.28 (d, *J* = 7.8 Hz, 2H, C$_8$**H**, C$_{22}$**H**), 7.68 (brd, *J* = 2.4 Hz, 2H, N$_{10}$**H**, N$_{14}$**H**), 3.87 (m, 2H, C$_{11}$**H**, C$_{15}$**H**), 2.70 (dd, *J* = 14.4 Hz, 4.2 Hz, 2H, C$_{12}$**H**, C$_{17}$**H**), 2.18 (dd, *J* = 14.4 Hz, 6.6 Hz, 2H, C$_{12}$**H**, C$_{17}$**H**)

$^{13}$C NMR (100 MHz, DMSO-*d*$_6$): δ124.4 (C**2**, C**19**), 108.7 (C**3**, C**18**), 127.3 (C**4,** C**26**), 118.3 (C**5**, C**25**), 118.5 (C**6**, C**25)**, 120.8 (C**7**, C**23**), 111.2 (C**8**, C**22**), 136.0 (C**9**, C**21**), 55.2 (C**11**, C**15**), 29.9 (C**12**, C**17**), 166.7 (C**13**, C**16**)

The absolution configuration of **4**, **7**, **10**, **13** and **16** were determined by comparing the specific rotation values of (**4** $[\alpha]_D^{22}$ -10 (*c* = 0.1, MeOH), **7** $[\alpha]_D^{22}$ -48 (*c* = 0.1, MeOH), **10** $[\alpha]_D^{22}$ -80 (*c* = 0.1, MeOH), **13** $[\alpha]_D^{22}$ -30 (*c* = 0.1, MeOH), **16** $[\alpha]_D^{22}$ -78 (*c* = 0.1, MeOH)) to reported values.[1]

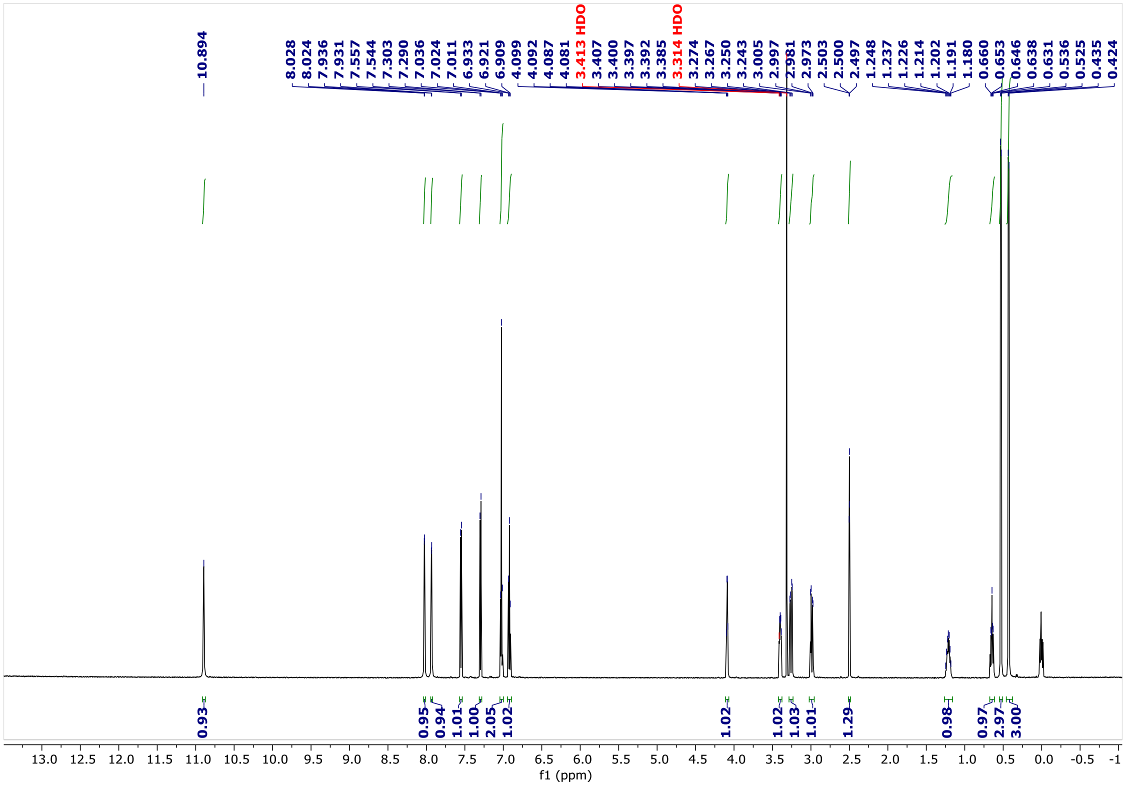



**Figure S1.** $^{1}$H NMR spectrum of **4** (DMSO-$d_6$, 600 MHz)

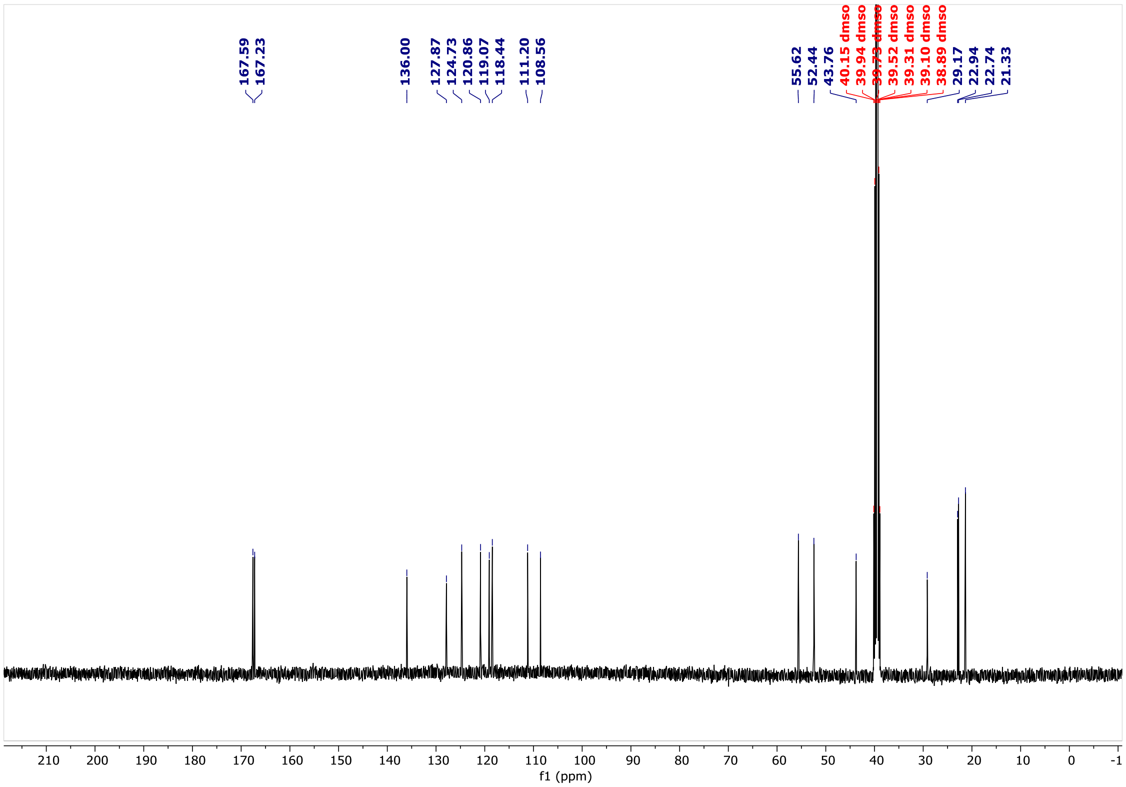



**Figure S2.** $^{13}$C NMR spectrum of **4** (DMSO-$d_6$, 100 MHz)

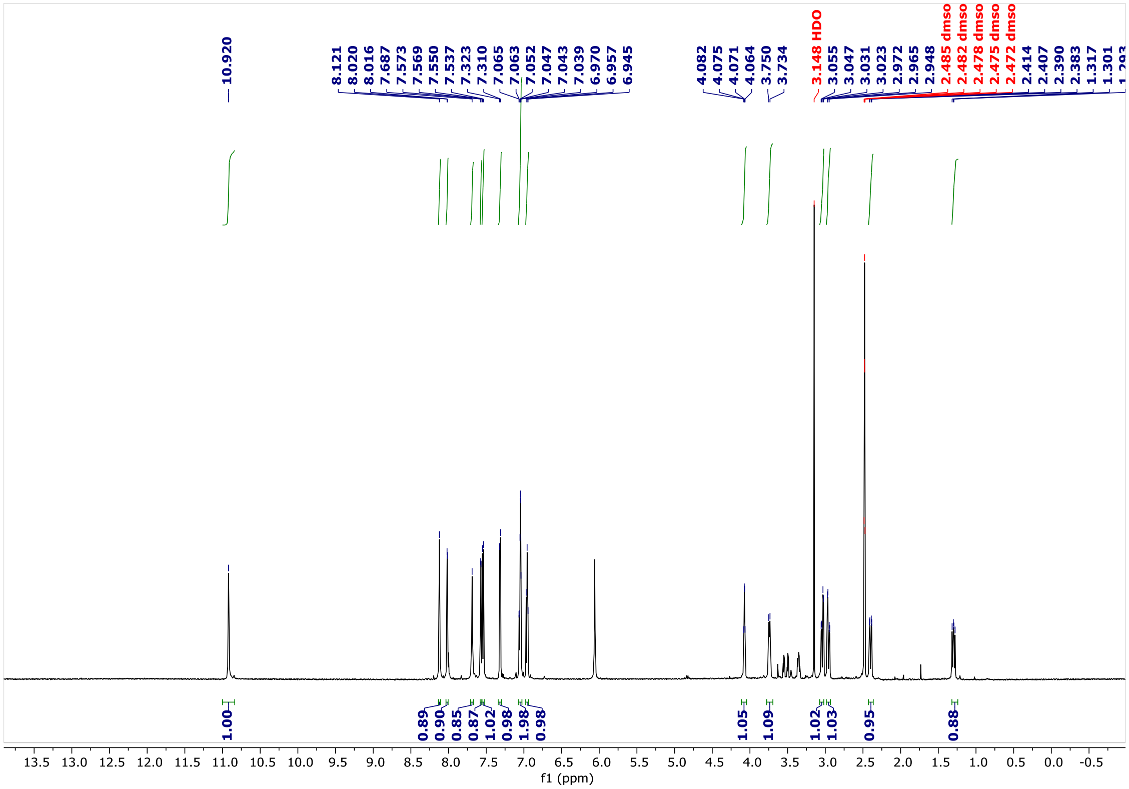


**Figure S3.** $^{1}$H NMR spectrum of **7** (DMSO-$d_6$, 600 MHz)

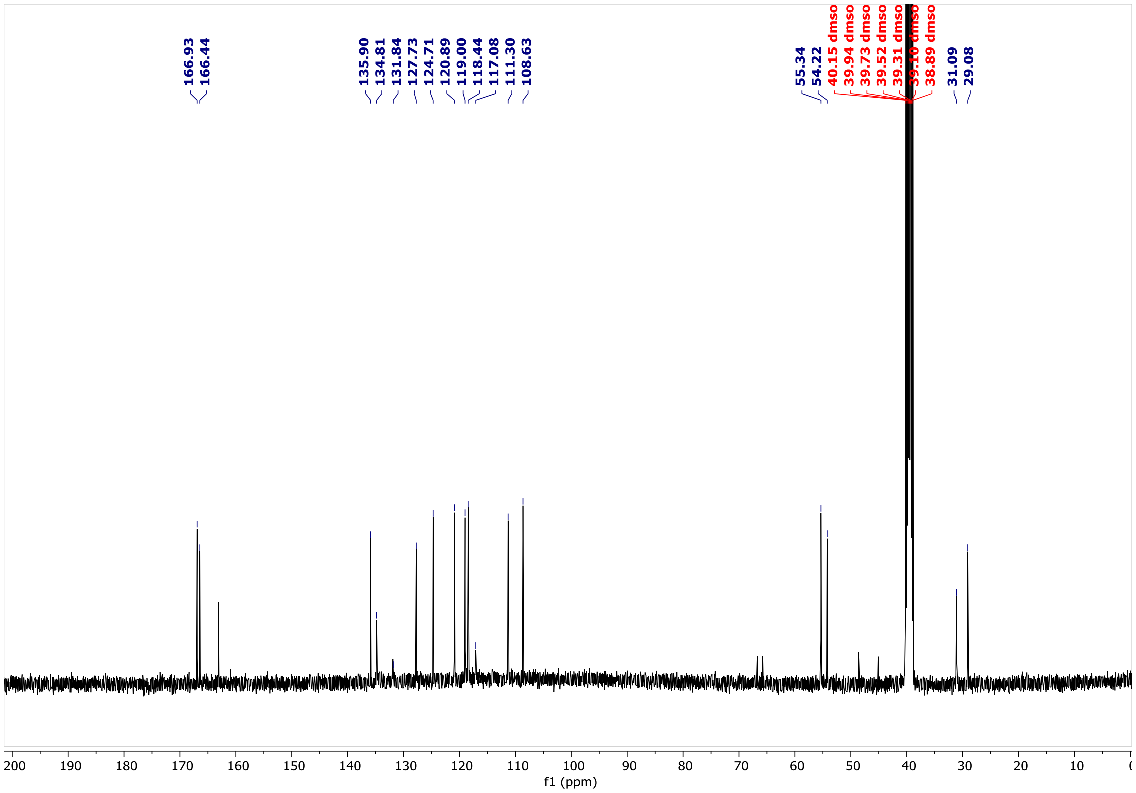


**Figure S4.** $^{13}$C NMR spectrum of **7** (DMSO-$d_6$, 100 MHz)

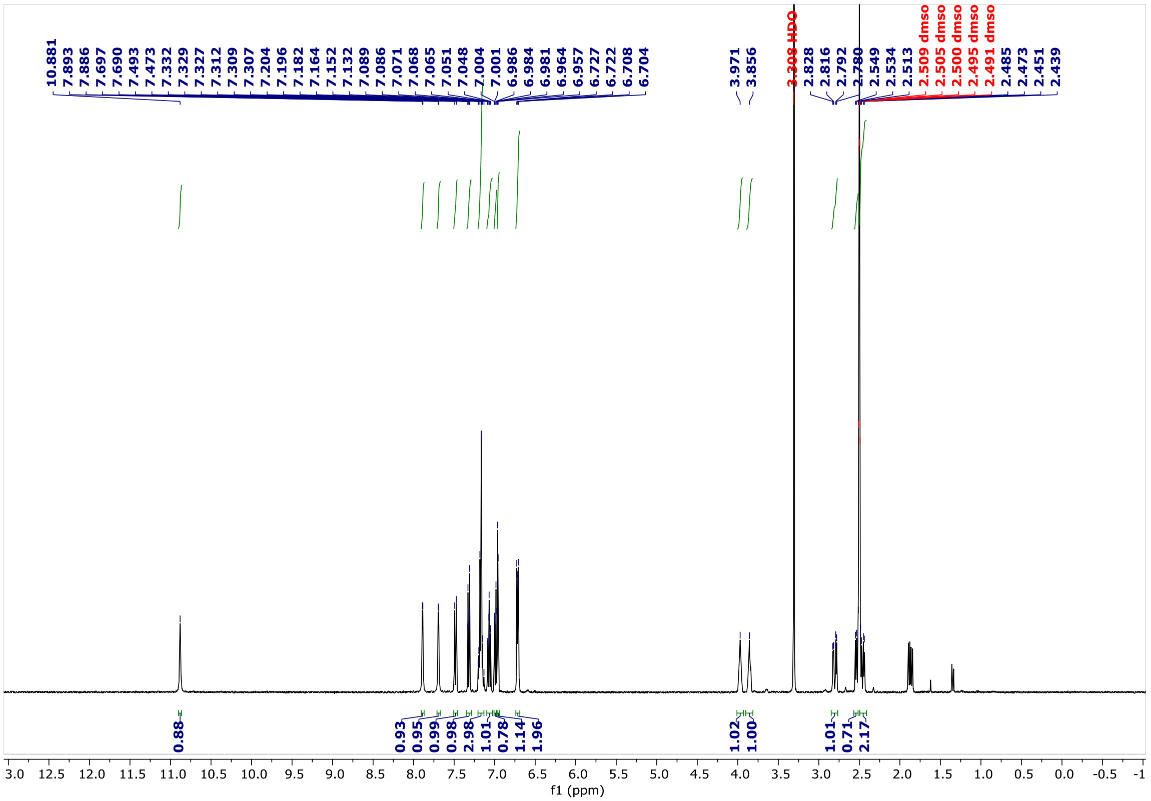


**Figure S5.** $^{1}H$ NMR spectrum of **10** (DMSO-$d_6$, 600 MHz)

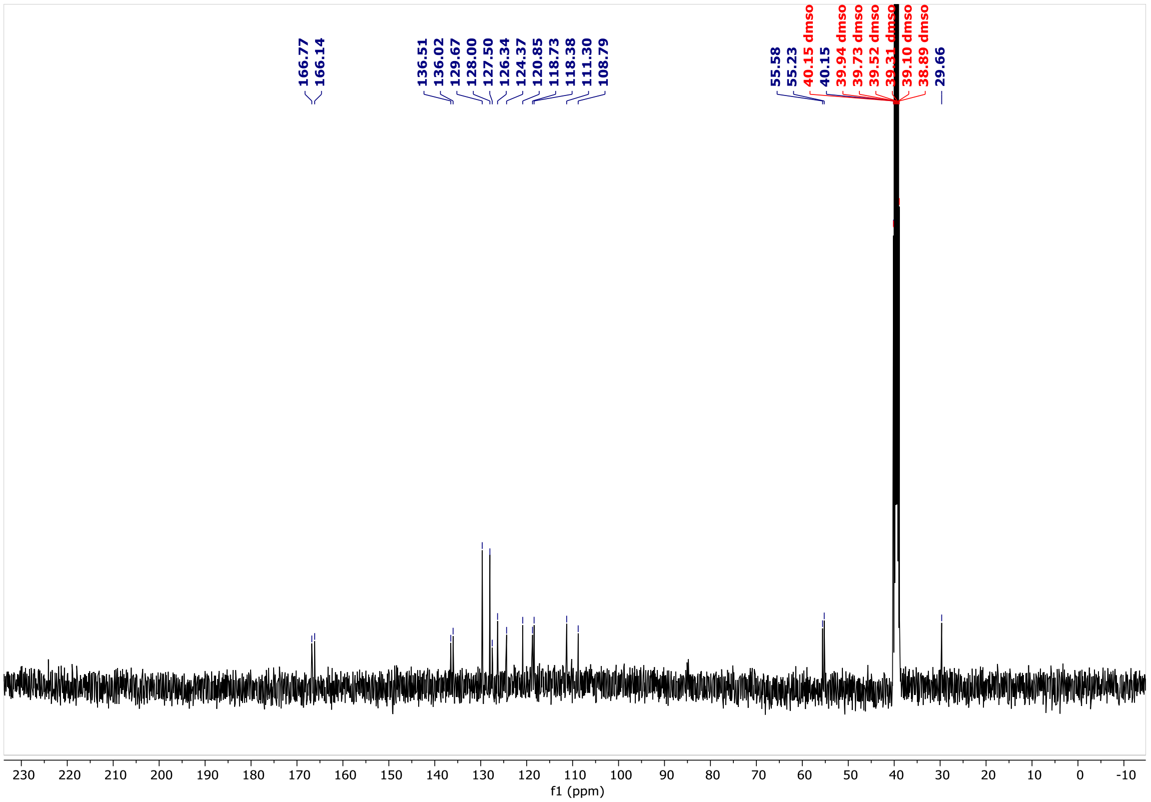


**Figure S6.** $^{13}C$ NMR spectrum of **10** (DMSO-$d_6$, 100 MHz)

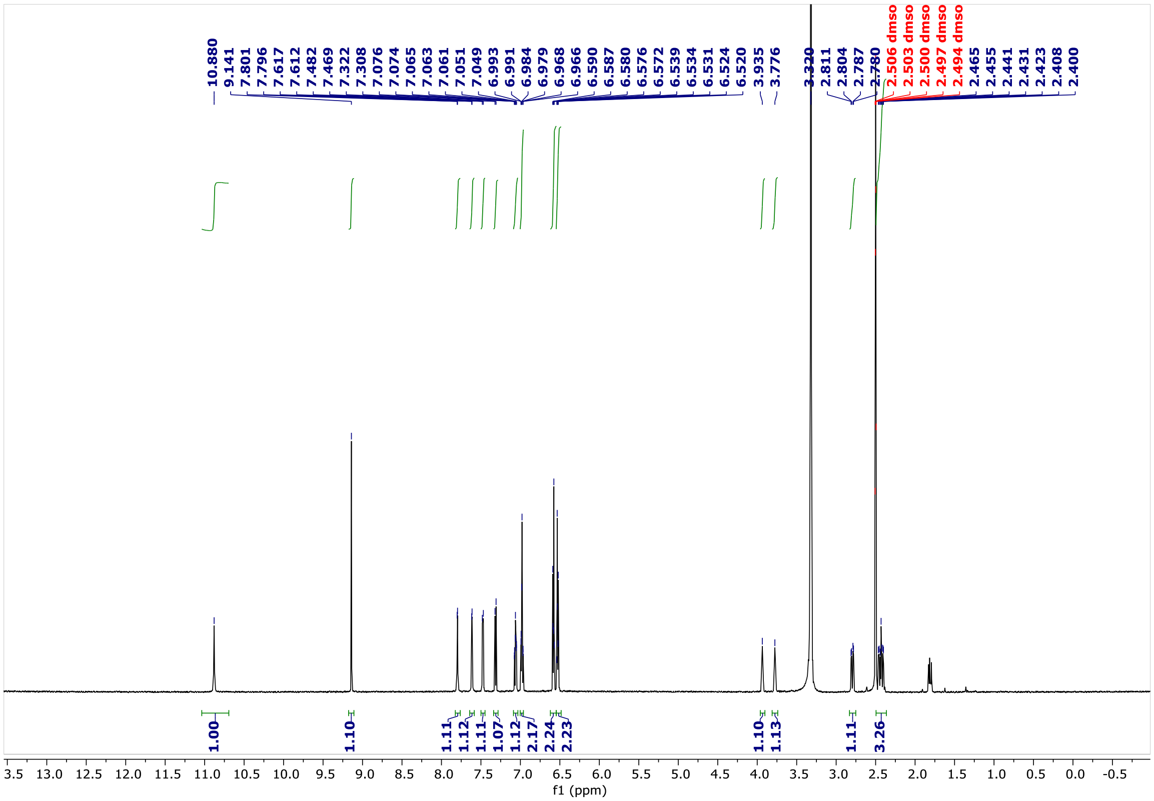


**Figure S7.** $^{1}$H NMR spectrum of **13** (DMSO-$d_6$, 600 MHz)

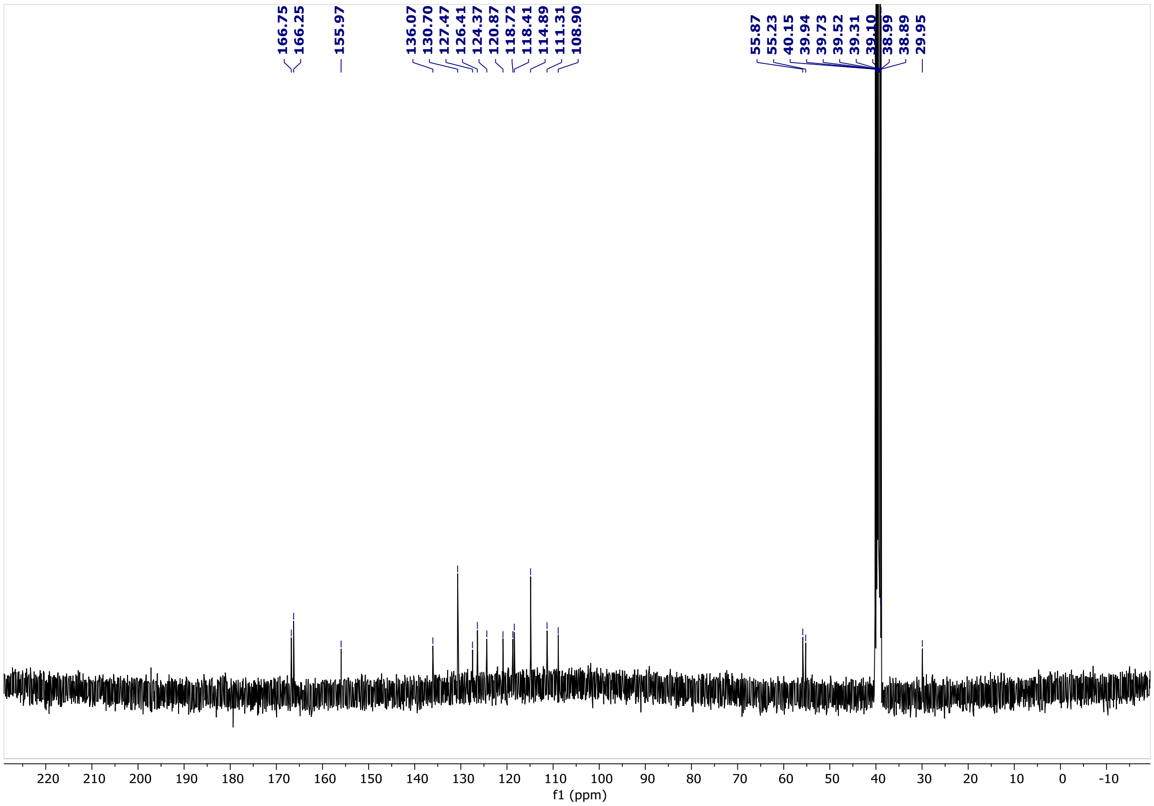


**Figure S8.** $^{13}$C NMR spectrum of **13** (DMSO-$d_6$, 100 MHz)

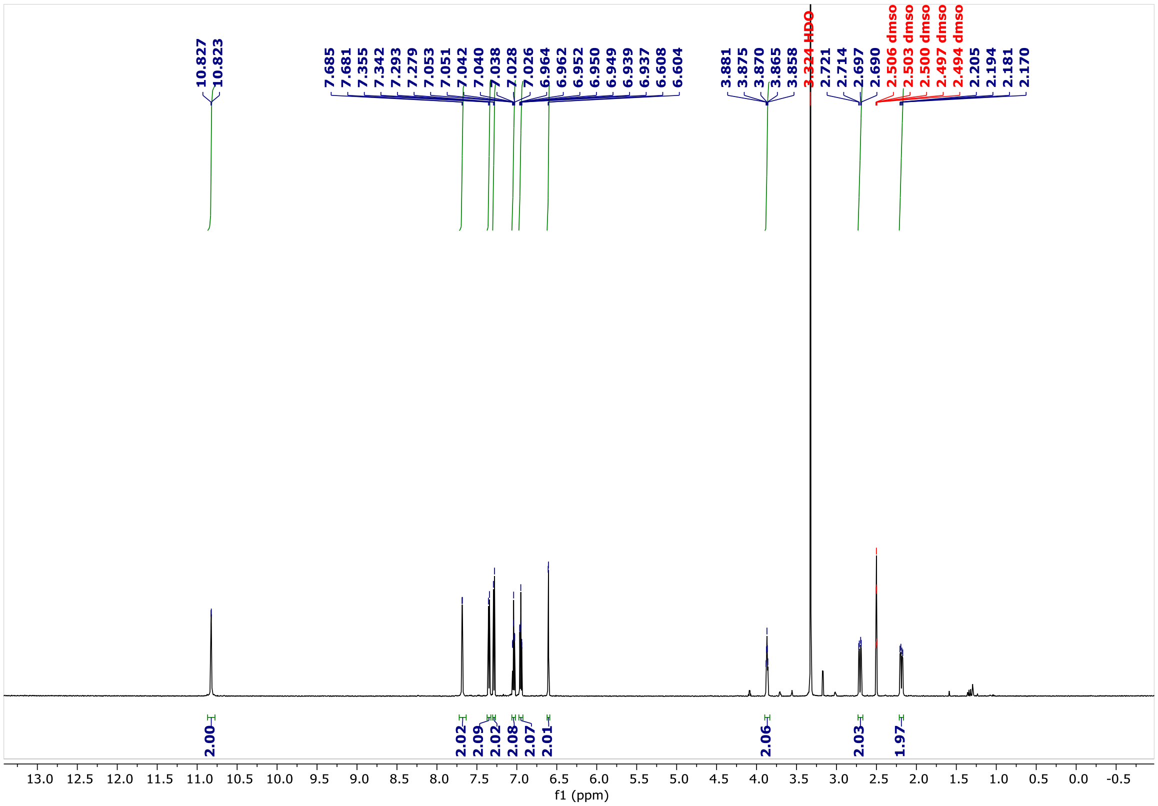


**Figure S9.** $^{1}$H NMR spectrum of **16** (DMSO-$d_6$, 600 MHz)

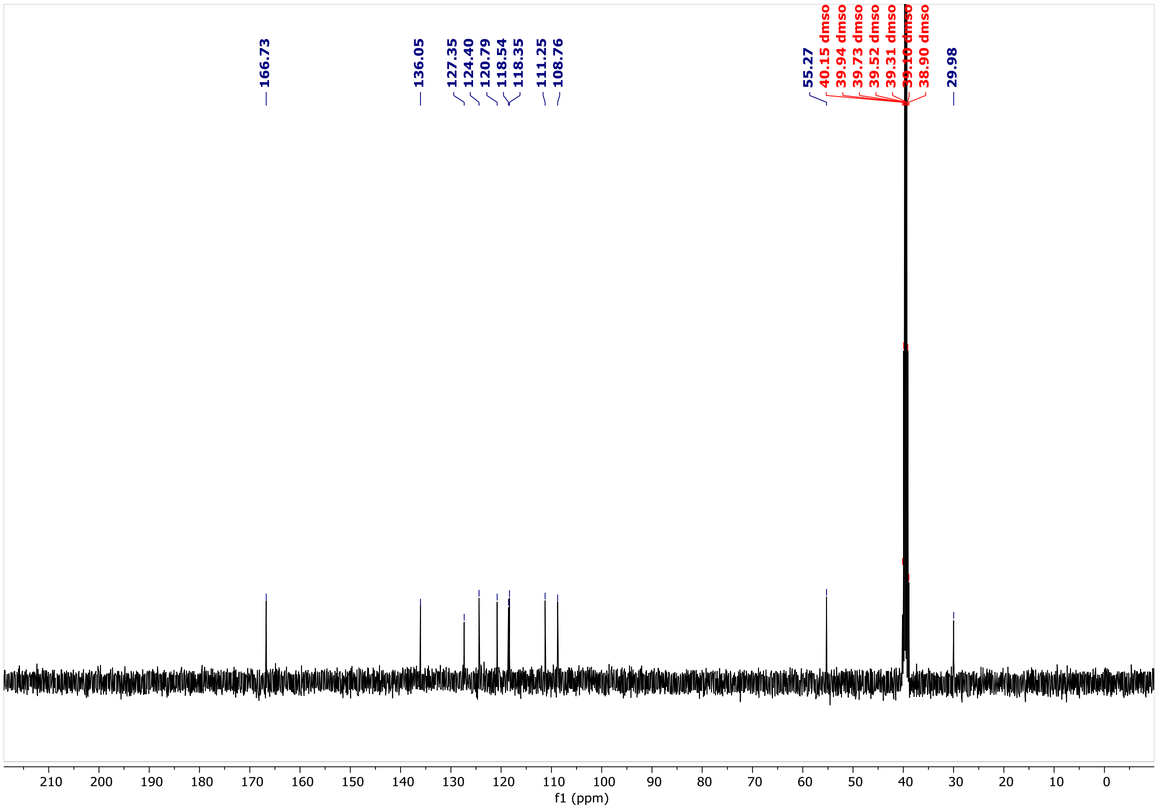


**Figure S10.** $^{13}$C NMR spectrum of **16** (DMSO-$d_6$, 100 MHz)
